# NERVE GROWTH FACTOR IS SUFFICIENT TO CAUSE MULTIPLE OSTEOARTHRITIS-RELEVANT PATHOLOGICAL FEATURES IN NAÏVE MURINE KNEE JOINTS

**DOI:** 10.1101/2025.06.20.660696

**Authors:** Alia M. Obeidat, Michael D. Newton, Jun Li, Baofeng Hu, Shingo Ishihara, Lindsey Lammlin, Lucas M. Junginger, Easton C. Farrell, Frank C. Ko, Richard J. Miller, Carla R. Scanzello, Tristan Maerz, Rachel E. Miller, Anne-Marie Malfait

**Author notes:** **Corresponding author:** Anne-Marie Malfait, MD PhD, Rush University Medical Center, 1611 W Harrison Street, Suite 510, Chicago IL 60612.

## Abstract

**BACKGROUND:** Nerve growth factor (NGF), a key mediator of pain and inflammation, is increased in joints with osteoarthritis (OA). Neutralizing NGF with monoclonal antibodies has shown analgesic effects in painful knee OA, but clinical development was stopped due to side effects in the joints. Knowledge about the biological effects of long-term exposure of joint tissues to NGF is limited. Therefore, we aimed to explore the effects of repeated intra-articular (IA) injections of NGF into the knee joints of healthy mice on pain and sensitization, as well as joint innervation and structure.

**METHODS:** We conducted five experiments in male C57BL/6 mice. In Experiment 1, NGF (50ng or 500ng) or vehicle was injected IA into the knee of naive wildtype (WT) mice, twice a week for 4 weeks. We assessed knee swelling, knee hyperalgesia and histopathology. In Experiment 2, mice were injected with 500ng NGF or vehicle, twice a week for 4 weeks and microCT of the knee was performed. In Experiment 3, Na_V_1.8-tdTomato reporter mice were injected with 500ng NGF or vehicle, twice a week for 4 weeks, and joint innervation was assessed. In Experiment 4, WT mice received 500ng NGF or vehicle twice a week for 4 weeks and were used for single cell RNA sequencing (scRNAseq) of the synovium. In Experiment 5, L3-L5 DRGs of mice that received 3 IA injections of 500ng NGF or vehicle twice a week were used for bulk RNA sequencing.

**RESULTS:** Repeated bi-weekly IA injections of NGF caused knee hyperalgesia in naïve mice. NGF caused dose-dependent knee swelling, synovial pathology, increased bone mineral density and trabecular bone thickness in the medial subchondral bone, growth of pre-osteophytes in the medial compartment, but no cartilage degeneration. NGF injection caused sprouting of Na_V_1.8+ neurons in the medial but not the lateral synovium. ScRNAseq of the synovium revealed upregulated genes related to neuronal sprouting, synovial fibrosis and ossification, confirming histopathological findings. Bulk RNA seq of DRG showed upregulated pathways related to axonal growth.

**CONCLUSIONS:** In healthy mouse knees, NGF induced mechanical sensitization, synovitis, neoinnervation in the medial synovium, subchondral bone changes and pre-osteophyte growth in the medial compartment, thus capturing many pathological changes observed in OA, except cartilage damage.

## INTRODUCTION

The neurotrophin, nerve growth factor (NGF), is essential for the growth and survival of sensory and sympathetic neurons during embryonic development (1). In adulthood, NGF is produced at low basal levels, but it is rapidly upregulated during inflammation to act as a key mediator of neuroimmune interactions in inflammatory processes and pain (2). NGF can be produced by many cell types, including epithelial and endothelial cells, fibroblasts, smooth muscle cells, glial cells, and immune cells such as lymphocytes, granulocytes, mast cells, eosinophils, and macrophages (reviewed in (2)). NGF exerts its effects through binding to two classes of receptors: the high affinity receptor, tyrosine receptor kinase A (TrkA) and the low affinity p75 neurotrophin receptor (p75R, p75^NTR^). Both receptors are broadly expressed and involved in neuronal development and pain sensitization (3,4).

NGF has been associated with several painful inflammatory conditions (2,4,5). In particular, NGF has received extensive attention as a target for chronic pain associated with osteoarthritis (OA) (6). OA is the most common form of arthritis, and with an estimated 600 million people globally suffering from symptomatic OA (7), it is one of the major sources of chronic pain in the world. Current pain management options include conventional analgesics such as non-steroidal anti-inflammatory drugs (NSAIDs) and intra-articular (IA) strategies such as hyaluronan or corticosteroids, but these show limited efficacy and are associated with safety concerns upon prolonged use (8). Clinical trials with neutralizing anti-NGF antibodies showed considerable efficacy in treating moderate to severe painful knee OA (9). However, drug development programs were ceased after the U.S. Food and Drug Administration (FDA) failed to approve anti-NGF antibodies for OA pain in 2021, owing to the risk of rapidly progressive OA (9). The mechanisms underlying this adverse effect are poorly understood, but the observation that sequestering NGF with antibodies may have adverse effects on joint integrity raises questions as to the biological function of NGF in the joint, and its putative role in maintaining joint homeostasis during disease. Indeed, both NGF receptors are widely distributed in joint tissues; for example, TrkA is not only expressed by nociceptors, but by a wide array of cells, including cholinergic neurons, satellite glial cells in the DRG, immune cells (macrophages, T cells, plasma cells, and mast cells), chondrocytes, and fibroblasts (4,10,11). The p75^NTR^ is expressed by DRG neurons, sympathetic ganglia, myeloid cells, satellite cells, Schwann cells, chondrocytes, fibroblasts, and immune cells (5,10–12).

Increased levels of NGF are characteristic of OA. NGF protein levels are elevated in OA synovial fluid, as they are in rheumatoid arthritis and spondyloarthritis (13,14) and in the serum of individuals with OA (14). In joints of people or experimental animals with OA, NGF mRNA is increased in chondrocytes (15–18), as well as in synovial fibroblasts and macrophages (19–21). The sensitizing and pain-producing effects of IA NGF injections into rodent knee joints have been well documented. A single IA injection into healthy rat knees induced weight-bearing asymmetry, mechanical allodynia in the hind paw, and knee swelling with synovial macrophage infiltration (22). In naïve mice, repeated IA injections of NGF resulted in knee swelling and knee hyperalgesia (23). In contrast, little is known about the effects of NGF on different joint tissues, especially not in a whole-joint *in vivo* context. In order to address this gap in knowledge and enable further development of analgesic strategies that target this pathway, we aimed to assess the biological actions of NGF in the whole murine knee. We sought to explore the effects of repeated IA injections of NGF into the knee joints of naïve, young adult mice on knee swelling and knee hyperalgesia, on histopathology and innervation of joint tissues, and on the molecular architecture of the synovium and the innervating dorsal root ganglia (DRG) in order to gain insight into how NGF may regulate processes involved in joint damage and pain.

## MATERIALS AND METHODS

### Animals

We used a total of n = 65 male C57BL/6 mice, either wildtype (WT) (inbred at Rush) or Na_V_1.8-tdTomato mice (which express a fluorescent tdTomato reporter in >90% of nociceptors) (24,25). Animals were housed with food and water *ad libitum* and kept on a 12-hour light cycle. All animal procedures were approved by the Institutional Animal Care and Use Committee at Rush University Medical Center.

### Injection protocols and experimental designs

All injection protocols are depicted in **Fig. 1**. In Experiment 1, NGF (R&D systems, # 1156-NG-100), 50ng or 500ng in 5 μL, or vehicle control (0.1% BSA) (05470, Sigma) (n=5 mice/group) were injected IA into the right knee joint of naive 12-week-old male WT mice, twice a week for 4 weeks. These mice were used to evaluate knee swelling and knee hyperalgesia, and to assess knee histopathology. In experiment 2, a cohort of Na_V_1.8-tdTomato mice was used to assess the medial and lateral subchondral bone of the tibia using micro-computed tomography (μCT) (n=4 vehicle and n=6 NGF, 500ng). This cohort had been used in a previous study from our group (26). In experiment 3, we used 12-week-old male Na_V_1.8-tdTomato mice and injected NGF (500ng in 5 μL) or vehicle control IA into the right knee joint (n=5 mice/group), twice a week for 4 weeks. These mice were used for innervation studies. In experiment 4, 12-week-old WT mice received NGF (500ng) or vehicle twice a week for 4 weeks (n=9/group), and mice were assessed for knee hyperalgesia. After 4 weeks of injections, synovium was collected for single cell RNA sequencing (scRNAseq) from 4 randomly selected mice. Lastly, in Experiment 5, 15-week-old WT mice received NGF (500ng) or vehicle twice a week for 10 days (3 IA injections in total, n=6/group). Twenty-four hours after the last injection, L3-L5 ipsilateral DRGs were collected for bulk RNAseq.

**Figure 1:**
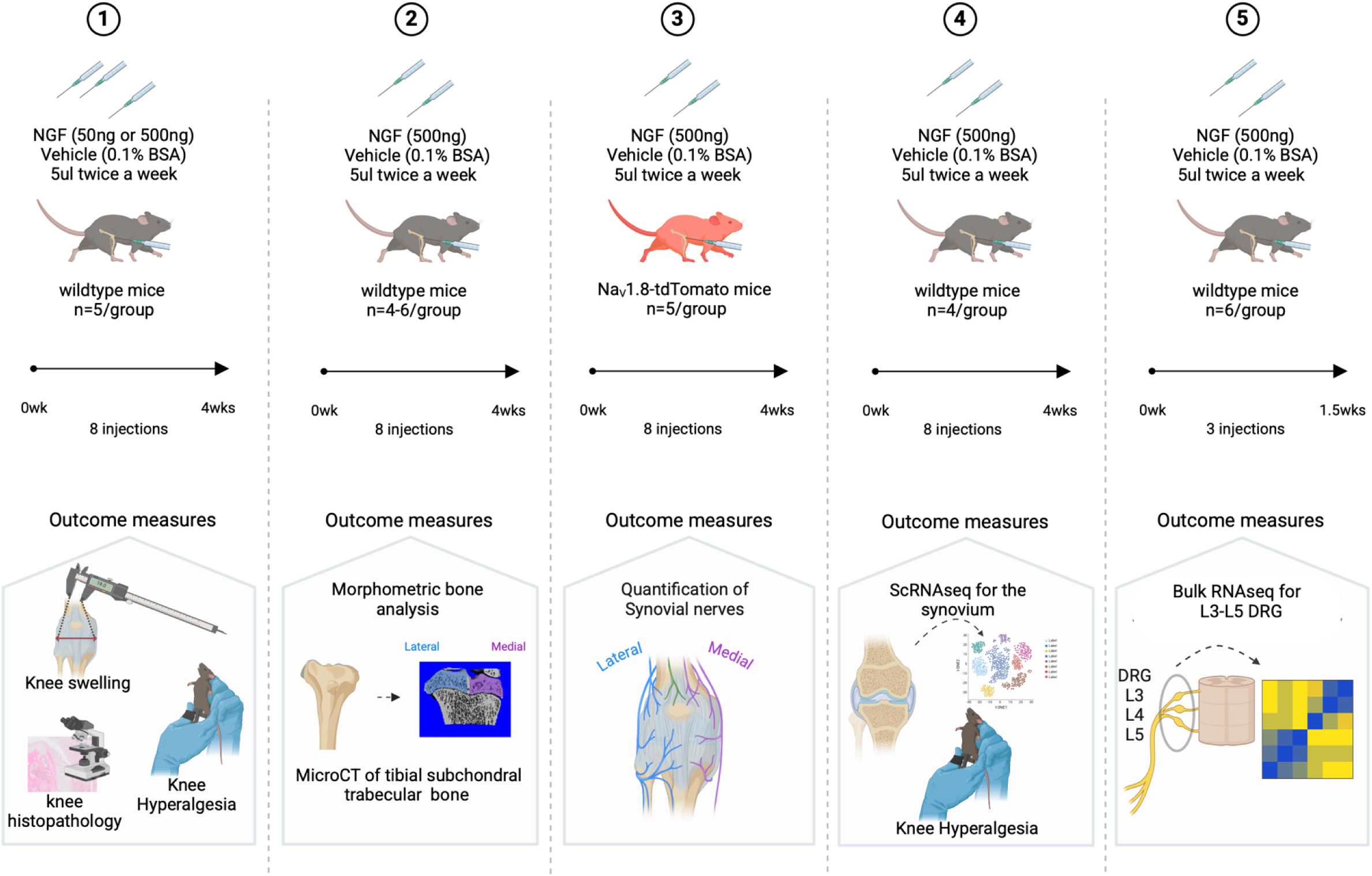
Schematic illustration of the 5 experiments performed, showing injection protocols, mouse strains (n), experimental endpoints, and outcome measures. ScRNAseq= single cell RNA sequencing. Created with BioRender.com.

### Knee swelling and knee hyperalgesia

In Experiment 1, knee width was measured at baseline (before the first injection) and then twice a week for 4 weeks, right before each injection, using micro-calipers as described (23). Knee hyperalgesia was assessed at baseline and then at weeks 2 and 4 by an observer blinded to treatment groups (SI). We used a Pressure Application Measurement (PAM) device, as described (27). Briefly, animals were restrained, and the PAM transducer was pressed against the medial side of the flexed ipsilateral knee. PAM software guided the user to apply a constantly increasing force (30 g/s) up to a maximum of 450 g, and the force at which the mouse tries to withdraw its knee was recorded. Two measurements were obtained per knee, and the withdrawal thresholds were averaged and reported.

### Knee Histopathology

Mice in Experiment 1 were euthanized after 4 weeks of injections (at the age of 16 weeks), and the right knees were collected. The knees were fixed in 10% natural buffered formalin, decalcified in EDTA for 3 weeks, embedded in paraffin, and sectioned at 5μm. Sections from the center of the joint were stained with hematoxylin (1x-Sigma HHS160) and eosin (1%-Sigma E4382) (H&E) for evaluation of cartilage damage based on modified Osteoarthritis Research Society International recommendations (28). For osteophyte scoring, one section with the major chondrophyte or pre-osteophyte was assessed using Osteomeasure software (OsteoMetrics), as described (29,30). Synovial pathology (synovial hyperplasia, cellularity, and fibrosis) was evaluated in four joint spaces (lateral femoral, medial femoral, lateral tibial, and medial tibial separately), as described (31). Five 5μm sections per mouse were assessed, 2 from anterior to the midpoint, a midpoint section, and 2 from the posterior area. Scoring was performed by 4 observers blinded to the treatment groups (AO, JL, BH, CRS). Scores were averaged for all five sections per area per mouse for each scorer. Scores from all 4 observers were averaged and graphed per synovial parameter.

### Micro Computed Tomography

Micro-computed tomographic (μCT) imaging was performed on WT mice, injected with either vehicle (n=4) or NGF (500ng, n=6) using a high-resolution laboratory imaging system (μCT50, Scanco Medical AG, Brüttisellen, Switzerland) in accordance with the American Society of Bone and Mineral Research (ASBMR) guidelines for the use of μCT in rodents (32). Scans were acquired using a 10 μm^3^ isotropic voxel, 70 kVp and 114 μA peak x-ray tube potential and intensity, 500 ms integration time, and were subjected to Gaussian filtration. The subchondral bone analysis was performed by manually contouring the outline of the medial and lateral tibial epiphysis separately. A lower threshold of 363 mg HA/cm3 was used to evaluate trabecular bone volume fraction (BV/TV, mm^3^/mm^3^), bone mineral density (BMD, mg HA/cm^3^), mean trabecular thickness (Tb.Th, mm), and mean trabecular separation (Tb.Sp, mm).

### Quantification of the neuronal signal in the synovium

For the innervation studies in Experiment 3, Na_V_1.8-tdTomato mice were perfused with 4% PFA at week 4, and right knees were collected. Twenty µm-thick coronal sections were collected from the mid-joint area as previously defined in (33). Sections were imaged using a laser-scanning confocal microscope (Olympus IX70) and processed using ImageJ and Fluoview software (FV10-ASW 4.2 Viewer). Adjustments were made to brightness and contrast to reflect true colors. All images were treated the same in terms of adjustments to brightness and contrast to minimize bias. Na_V_1.8+ signal was quantified in the medial and lateral synovium in mid-joint coronal sections using Image J (30). Briefly, regions of interest (ROI) were manually outlined for each section using anatomical landmarks. Thresholds were adjusted for all images similarly to control for background. The area of positive signal within each ROI was measured and normalized to the total area of the ROI; the percentage of positive signal per ROI is reported. Quantification was performed by an observer blinded to the treatment groups. Na_V_1.8+ subchondral bone channels were quantified as previously described (33).

### Single cell RNA sequencing of the synovium

For single cell sequencing of the synovium, 8 C57Bl/6 mice (*n* =2/sex/condition, age 12 weeks) were randomized to receive 4 weeks of biweekly intra-articular injections of NGF (500 ng) or vehicle control. At the end of week 4, mice were humanely euthanized via CO2 asphyxia, and ipsilateral synovia were collected and digested to yield a single cell suspension, as described (34). Cell suspensions were frozen in liquid nitrogen until the time of submission. Single cell suspensions were submitted to the University of Michigan Advanced Genomics Core for barcoding and library prep via the 10X Genomics processing pipeline (Chromium Next GEM Single Cell 3’ Kit v3.1). Two bioreplicates were submitted per condition, with each bioreplicate pooled from one male and one female ipsilateral synovium. Pooled libraries underwent paired-end sequencing (100bp+100bp) on the DNBseq G400 system (MGI Tech). Raw data were aligned to the mm10-2020-A mouse transcriptome using STAR (35) and were preprocessed in Cell Ranger (v7.0.1). Quality control of sequencing indicated high accuracy across all samples (**Suppl. Fig. 1A)**.

Detailed methods for scRNAseq data analysis are provided in Suppl. Methods. Briefly, count matrices were generated using Cell Ranger and exported to R for analysis. Count matrices for individual bioreps were pre-processed using SoupX (36) to remove ambient RNA contamination and processed via the Seurat toolbox (37). Seurat objects were preprocessed, including pre-filtered to exclude debris, multiplets, and non-viable cells, further multiplet removal via DoubletFinder (38) (**Suppl. Fig. 1B**) and normalization and scaling using SCTransform (39). Samples were integrated into a single Seurat object and clustered to identify unique cell populations (Figure 5A-C). Clusters were annotated by expression of unique, cell type-defining gene markers. Spearman correlation confirmed high correlation between bioreps within each condition (**Suppl. Fig. 1C**). Within each cell type, differential expression (DE) analysis between NGF and Veh was performed to calculate NGF-induced differential gene expression. Functional annotation of gene-level results was performed within each cell type via gene set enrichment analysis (GSEA) via the *clusterProfiler* (40) implementation of the *fgsea* algorithm (41). Transcription factor binding motif (i.e. regulon) analysis was performed via RcisTarget (42), to determine which transcription factors were predicted to underpin expression of NGF-induced changes to synovial lining fibroblasts.

### Bulk RNAseq of lumbar DRGs

Ipsilateral DRGs (L3-L5) were collected from male WT mice (age 15 weeks) that received 3 IA injections of 500ng NGF (n=6) or vehicle (n=6) in 5μL over 10 days. DRGs were collected 24 h after the last injection and lysed in Trizol. RNA was extracted using an RNeasy micro kit (Qiagen) and sent to LC Sciences for sequencing. RNA Integrity Numbers all passed QC and were greater than 8.6. A poly(A) RNA sequencing library was prepared following Illumina’s TruSeq-stranded-mRNA sample preparation protocol. Paired-ended sequencing (150 bp) was performed on Illumina’s NovaSeq 6000 sequencing system (LC Sciences). HISAT2 was used to map reads to the Mus musculus reference genome (ftp://ftp.ensembl.org/pub/release-101/fasta/mus_musculus/dna/) (43). The mapped reads of each sample were assembled using StringTie with default parameters, and StringTie and ballgown were used to estimate the expression levels of all transcripts and perform expression abundance for mRNAs by calculating FPKM (fragment per kilobase of transcript per million mapped reads) value. The raw sequence data and counts matrices have been submitted to NCBI Gene Expression Omnibus (GEO) with accession number GSE287526. mRNA differential gene expression analysis was performed by R package DESeq2 (44) between two different groups (NGF vs. vehicle). The EnhancedVolcano R package was used to create the Volcano plot. Pathway analysis was performed by inputting upregulated or downregulated genes (padj < 0.05) into Cytoscape v3 and using the largest subnetwork of genes for STRING enrichment analysis for Reactome pathways with a redundancy cutoff of 0.5 applied. Transcription factor analysis was performed using the online ChEA3 tool (45); the mean score setting was used.

### Statistical analysis

Sample size was determined based on our previous data comparing innervation changes between sham and DMM (33). Unpaired two-tailed Student’s t test was used for pairwise comparisons of joint innervation and bone morphometric analyses. Ordinary two-way ANOVA followed by Sidak’s post-test was used for multiple group comparison of knee swelling and knee histopathology. Ordinary two-way ANOVA was used followed by Dunnett’s post-test for multiple group comparison of knee hyperalgesia. Statistical analyses were performed using GraphPad Prism 9. Data are presented as mean±SEM.

## RESULTS

### Intra-articular administration of NGF causes synovitis, accompanied by knee swelling and knee hyperalgesia

We first sought to assess the effects of repeated IA injections of NGF on the development of synovitis, knee swelling, and knee hyperalgesia (Experiment 1). We confirmed previous findings that repeated IA administration of NGF causes knee swelling (23). We tested NGF at two doses, 50 ng and 500 ng. The high dose (8 injections of 500 ng over 4 weeks) induced significant knee swelling as early as 4 days after the first injection, with a sustained effect up to 31 days. The low dose (8 injections of 50ng over 4 weeks) did not induce significant knee swelling (p=0.3011) (**Suppl. Fig. 2A,B).** We also measured knee hyperalgesia and found that all mice that received NGF, either 50ng or 500ng, developed knee hyperalgesia after 2 weeks of injections. By week 4, knee hyperalgesia was present in the high-dose group but had subsided in the low-dose group **(Suppl. Fig. 2C).**

Synovial histopathology was evaluated in all three groups after 4 weeks of injections, revealing striking effects of NGF **(Fig. 2)**. The high dose caused marked changes in both the medial and the lateral synovium, with an increase in subintimal cellularity, fibrosis, and lining hyperplasia compared to the vehicle group **(Fig. 2A-C)**. In mice that received the low dose of NGF, the effect was less pronounced, with a significant increase in cellularity in the medial synovium **(Fig. 2A)**, and a slight increase in fibrosis and hyperplasia that was not significant compared to vehicle controls (p= 0.0660 and p= 0.0856, respectively) **(Fig. 2B,C),** while there were no differences observed between the low NGF dose and vehicle groups in the lateral synovium **(Fig. 2A-C)**. Representative histological images of the pathological changes in the medial and lateral synovia are shown in **(Fig. 2D-F)**.

**Figure 2:**
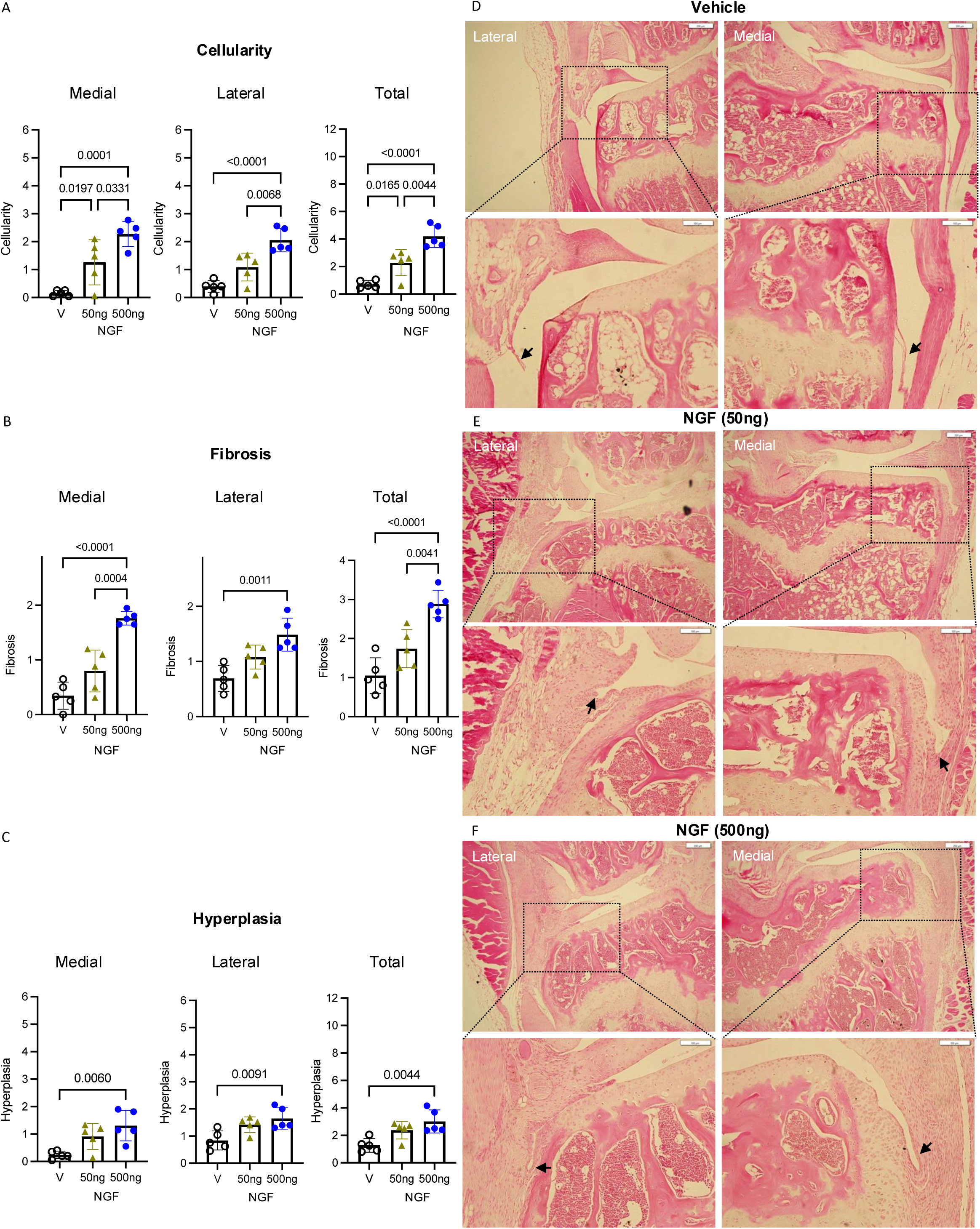
(A-C) Synovial cellularity, fibrosis and hyperplasia were evaluated in wild-type (WT) mice that received repeated intra-articular injections of recombinant murine NGF (IA 50 ng or 500 ng, 5 µL, 2x/week) or vehicle (5 µL, 2x/week) for 4 weeks; n=5 mice/group. Scores are shown for the medial and the lateral compartments, as well as the total joint scores; (D-F) Representative histological images of medial and lateral tibial synovia showing overview and zoomed in images (in black insets) from knee sections of mice injected with (D) vehicle, (E) 50 ng NGF, or (F) 500 ng NGF mice, respectively. Histological images show all 3 pathological features scored in (A-C). The zoomed in images in black insets show fibrosis and increased cellularity in NGF-injected mice. Black arrows show representative areas used to score hyperplasia. Two-way ANOVA with Sidak’s post-test. Scale bar = 200µm for overview images and 100µm for zoomed in images.

### Intra-articular administration of NGF causes subchondral bone changes but not cartilage damage

Chondrocytes express both NGF receptors, and cartilage from human OA patients and preclinical OA models exhibit increased NGF mRNA (11,15). We histologically scored cartilage damage but did not observe overt cartilage degeneration in any of the experimental groups after 4 weeks of injections (medial + lateral cartilage degeneration score, max score = 60) **(Fig. 3A)**. Articular surfaces were intact and congruent, and we observed no evidence of surface fibrillation characteristic of early OA. However, in the medial compartment of the knee, small chondrophytes or pre-osteophytes developed in response to NGF, and this occurred in a dose-dependent manner **(Fig. 3B)**. Representative histological images of the medial and lateral compartments are shown in **(Fig. 3B)**. Moreover, µCT analysis revealed that 8 injections of 500 ng NGF led to bone changes in the medial tibial subchondral bone, where bone mineral density and trabecular bone thickness were significantly increased compared to control knees, while no significant differences in bone volume fraction or mean trabecular spacing were observed between groups **(Fig. 3C-F)**. No changes in any of the bone morphometric parameters were observed in the subchondral bone of the lateral compartment compared to the vehicle group **(Fig. 3C-F)**. Representative 3D reconstructions of the subchondral trabecular bone from NGF-treated mice demonstrate a sclerotic morphology reminiscent of that observed in human and experimental OA (**Fig. 3G)**.

**Figure 3:**
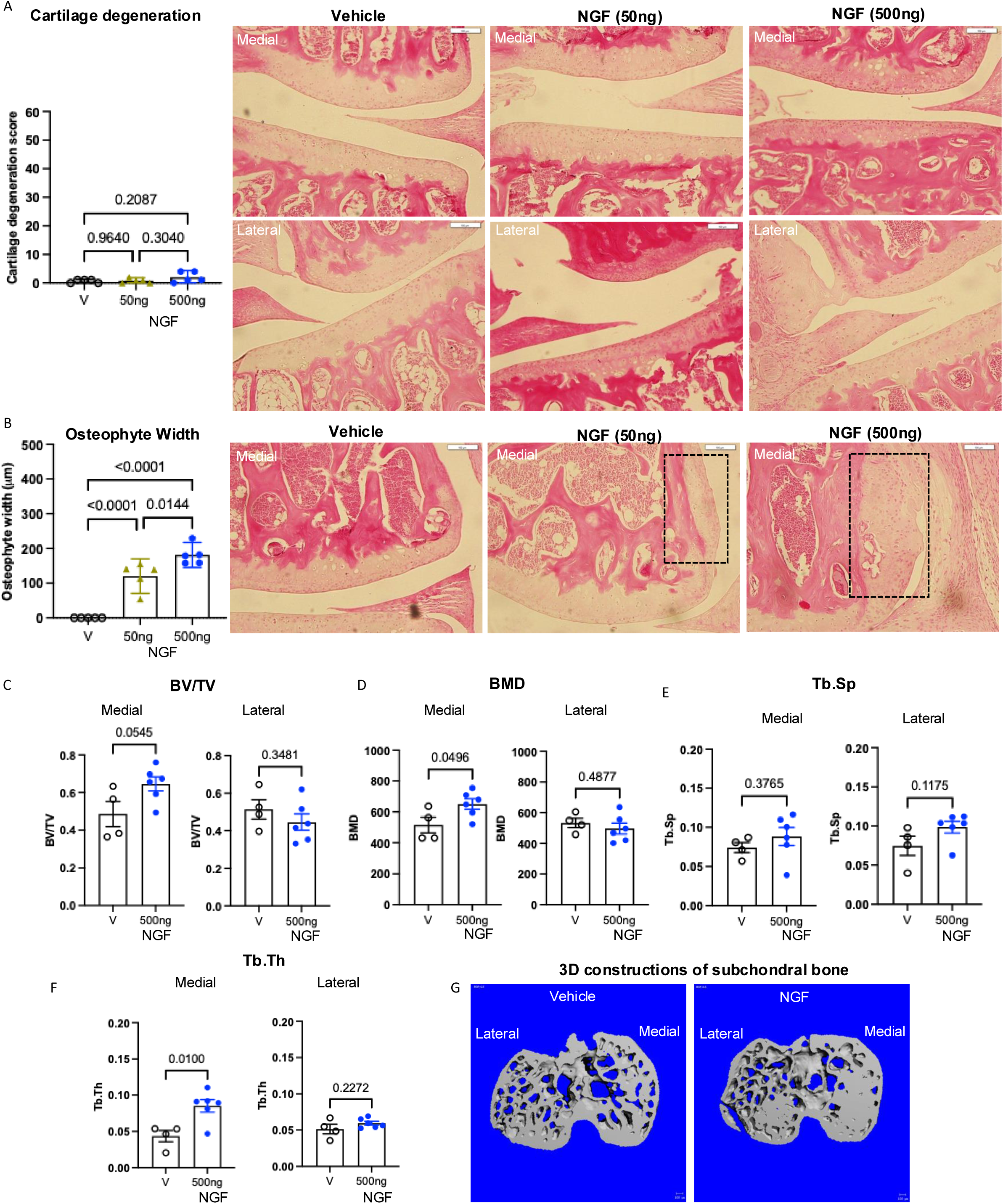
(A) Cartilage degeneration scores and representative histological images of knee sections from WT mice injected with vehicle, low NGF or high NGF, respectively; (B) Small chondrophytes or pre-osteophytes developed on the medial side. The graph shows the width in µm with representative histological images of wild-type (WT) mice injected with vehicle (5 µL, 2x/week), low NGF (50 ng, 5 µL, 2x/week), or high NGF (500 ng, 5 µL, 2x/week) for 4 weeks, n=5 mice/group. Black insets in B showing examples of developing pre-osteophytes in low dose and high dose NGF groups; (C-F) Bone volume / Trabecular volume (BV/TV) ratio, bone mineral density (BMD, in mm), mean trabecular thickness (Tb.Th, in mm), mean trabecular separation (Tb.Sp, in mm) respectively in the medial and lateral tibial subchondral trabecular bone from knees injected with NGF (500ng, n=6) or Vehicle (n=4); (G) Representative µCT scans of the subchondral trabecular bone from vehicle and NGF groups, respectively. For A and B, we used two-way ANOVA with Sidak’s post-test. For C-F, we used unpaired two-tailed t test. Scale bar=100µm.

### Intra-articular injection of NGF causes nociceptor sprouting in the medial synovium and into growing pre-osteophytes

Nociceptor sprouting is observed in subchondral bone and in synovium during progression of experimental OA (33,30). To determine whether NGF by itself is sufficient to induce this process thought to underpin pain in OA, we evaluated the Na_V_1.8+ nociceptive innervation of the knees after 8 injections of 500ng NGF or vehicle. We chose to assess knee innervation in mice that received the high dose of NGF, as we had observed more prominent pathological changes in this group in Experiment 1. In the medial synovium, we observed no or very few nociceptive fibers in the vehicle-injected group, which is in line with what we previously reported in naïve young mice (33), whereas Na_V_1.8+ fibers had sprouted in the deep layers of the medial synovium of NGF-injected knees **(Fig. 4A-C).** Representative confocal images of the medial synovium in NGF and vehicle treated groups are shown in (**Fig. 4B,C**). The naïve lateral synovium of these 16-week old mice contained more nociceptors than the medial synovium, as we have reported before (33), and no significant changes were detected among the treatment groups **(Fig. 4D)**. Representative confocal images of the lateral synovium in NGF and vehicle treated groups are shown in **(Fig. 4E,F)**. We did not detect changes in the innervation pattern of the medial and the lateral subchondral bone between the two groups (**Suppl. Fig. 3A-C)**, but we observed Na_V_1.8+ nociceptors sprouting toward growing pre-osteophytes; a representative image is shown in (**Suppl. Fig. 3D**).

**Figure 4:**
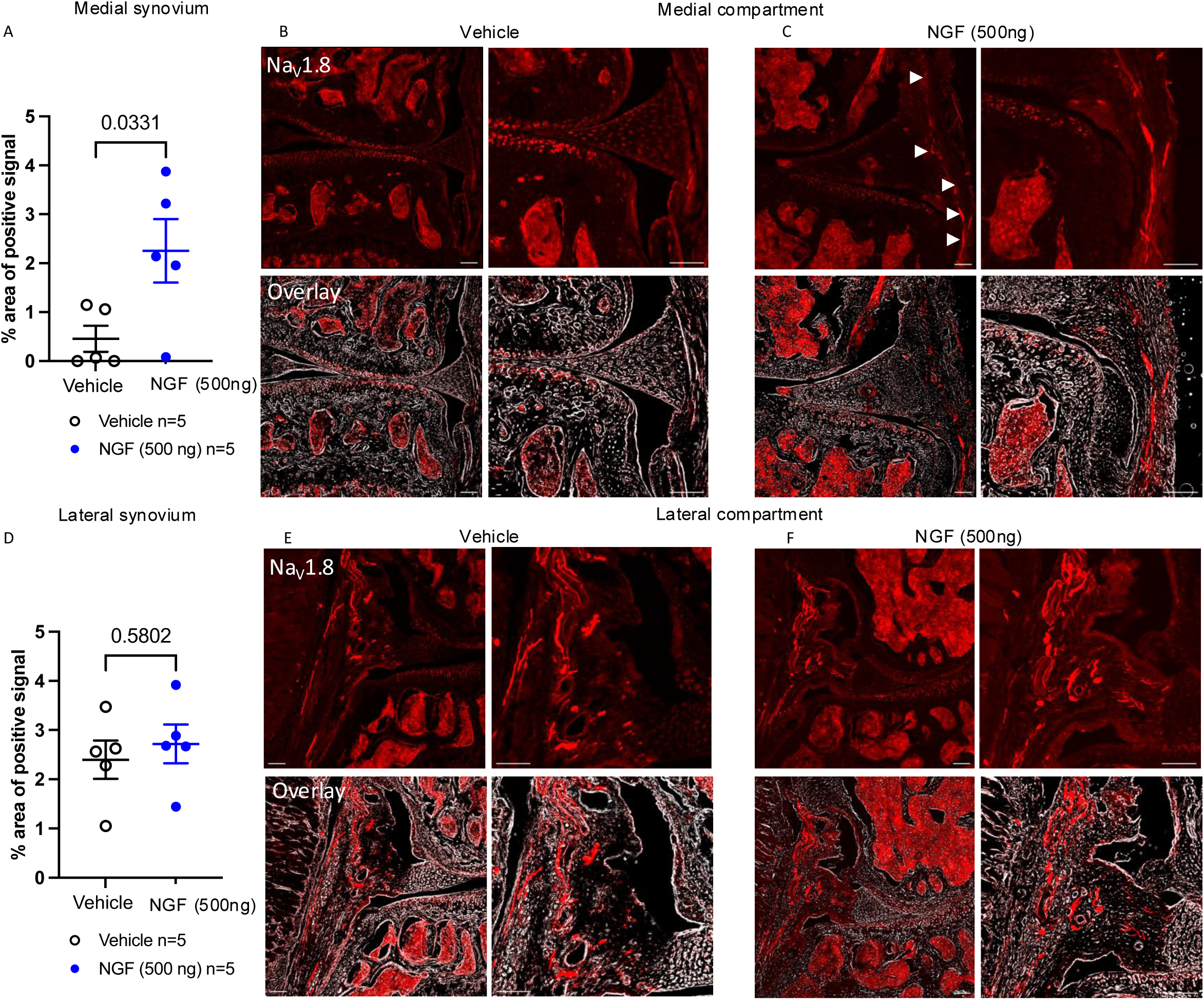
(A) Quantification of neuronal signal in the medial synovium of Na_V_1.8-tdTomato mice injected with vehicle (5 µL, 2x/week) or recombinant murine NGF (IA 500 ng, 5 µL, 2x/week) for 4 weeks; n=5 mice/group; (B,C) Representative confocal images of the medial compartment of vehicle and NGF treated groups respectively showing Na_V_1.8 fibers in white arrows; (D) Quantification of neuronal signal in the lateral synovium of Na_V_1.8-tdTomato mice injected with vehicle (5 µL, 2x/week) or recombinant murine NGF (500 ng, 5 µL, 2x/week) for 4 weeks; n=5 mice/group; (E,F) Representative confocal images of the lateral compartment of vehicle and NGF treated groups, respectively. Scale bar=100µm.

### Single cell RNA sequencing of the synovium

Given the profound pathological changes we observed in synovium following NGF administration, we next sought to assess which cell types and transcriptional signatures underpin this effect. To achieve this in a cell type-specific manner, we performed scRNAseq on synovium from mice injected IA with either NGF or vehicle. Consistent with Experiment 1, we confirmed that mice in this experiment also developed knee hyperalgesia after 4 weeks of injections of the high dose **(Suppl. Fig 4).**

Following quality control and preprocessing, a total of 17,041 high quality cells were identified, with relatively equal contributions from Veh (n=8,170) and NGF (n=8,871) conditions (**Suppl. Fig. 5A)**. Twelve major cell types were identified and annotated **(Fig. 5A)**, which were consistent with our previous experience in the murine synovium. All cell types were transcriptionally distinct **(Suppl. Fig. 5E)** and expressed canonical cell markers consistent with their lineage **(Fig. 5C, Suppl. Fig. 5D)** (34). NGF-treated samples had marginally higher sublining fibroblast content **(Fig. 5B)**, consistent with our observation of synovial fibrosis **(Fig. 2A-C),** but all other cell types did not vary markedly in abundance between the two groups. We did not observe major changes in relative cellular abundances or the emergence of any transcriptionally-distinct cell types.

**Figure 5:**
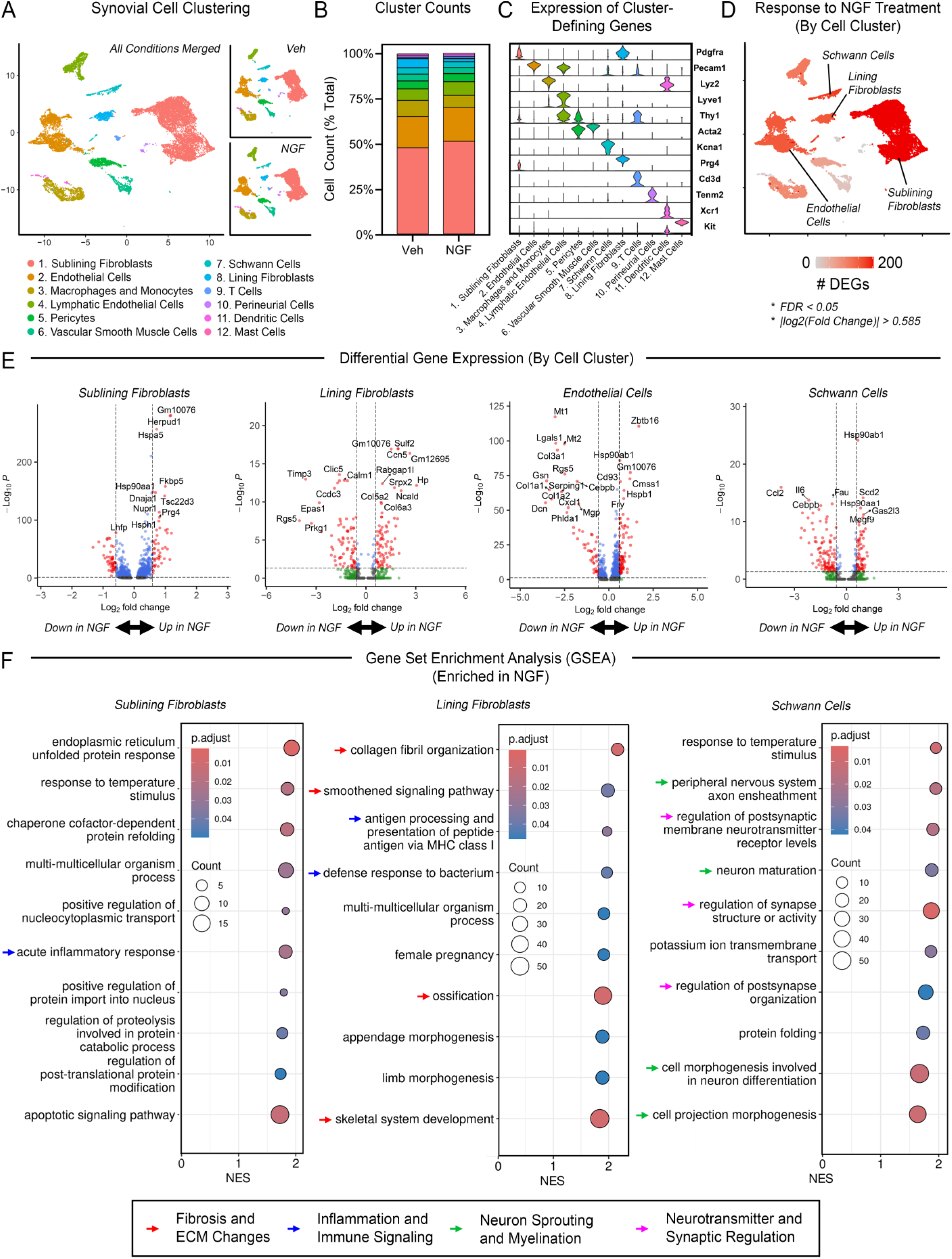
Single cell RNA sequencing of whole synovium response to intra-articular NGF injection. (A) Annotation of distinct synovial cell clusters. Twelve major cell types were identified and annotated (fibroblasts (sublining and lining), endothelial cells (vascular and lymphatic), and macrophages/monocytes comprised the majority, and smaller populations of pericytes, vascular smooth muscle cells (VSMCs), Schwann cells, T cells, perineurial cells, and dendritic cells were also detected; (B) Proportional cell counts of synovial cells in Veh and NGF groups were similar; (C) Expression of cluster-defining genes; (D) Cluster-specific response to NGF treatment, enumerated as # DEGs per cluster, shows the largest number of NGF-induces differences in gene expression occurred in sublining and lining fibroblasts, endothelial cells, and Schwann cells. NGF-induced differential gene expression of these clusters is shown as (E) volcano plots of differential gene expression (cutoffs at FDR < 0.05 and |log2(fold change)| > 0.585, and (F) gene set enrichment (top 10 enriched pathways shown, sorted by normalized enrichment score, NES).

We performed DE analysis within each cluster to assess which cell types were most perturbed by NGF treatment, along with gene-set enrichment analysis to functionally annotate the signaling pathways activated by NGF treatment. NGF induced a robust response across a broad range of cell types, most notably lining and sublining fibroblasts (lining: 154 DEGs; sublining: 102 DEGs), vascular endothelial cells (154 DEGs), and Schwann cells (180 DEGs) (*P_adj_* < 0.05, |log_2_(fold change)| > 0.585, **Fig. 5D-E**, **Suppl. Fig. 6-7**). Pathway analysis demonstrated that vascular endothelial cells were enriched in pathways consistent with the structural changes observed in NGF-treated knees **(Fig. 5F)**. Schwann cells were enriched in multiple pathways related to neuronal growth and myelination, marked by upregulation of genes including *Megf9* and *Gas2I3* (46). Schwann cells also upregulated multiple heat shock protein pathway genes (*e.g.*, *Hsp90aa1*, *Hsp90ab1*) which have been postulated to play a role in neuron development and regulation (47). Activation of heat shock protein pathways was also observed across other cell types, including sublining fibroblasts, vascular and lymphatic endothelial cells, macrophages and monocytes, pericytes, and VSMCs, suggesting that NGF may activate a broad stress response mechanism. Sublining fibroblasts and lining fibroblasts both exhibited enrichment of pro-inflammatory pathways **(Fig. 5F)**. Lining fibroblasts were enriched in pathways related to innate immunity, marked by upregulation of *Hp*, the gene encoding haptoglobin, shown to be involved in mediating innate immune responses (48,49), as well as histocompatibility genes (*e.g., H2-D1*, *H2-K1*, *H2-T22*). Lining fibroblasts in particular exhibited NGF-induced enrichment of pathways related to the histopathological effects observed in synovium, including fibrosis (collagen fibril organization, smoothened signaling pathway) and osteochondral bone formation (ossification, skeletal system development) **(Fig. 5F)**.

Gene-level analyses revealed that lining fibroblasts increased their expression of multiple collagens, including fibrosis-associated types 3, 5, and 6 (*Col3a1*, *Col5a1*, *Col6a1*), crosslinking enzymes (*e.g., Lox*, *Loxl1*), and linking glycoproteins (e.g. Emilin1) along with increased expression of genes related to fibrosis-associated signaling pathways including Wnt (*e.g.*, *Rspo2*, *Serpinh1*), and Smoothened (e.g. *Smo*, *Gli3*, *Gas1*) (50–53) **(Fig. 6A)**. Lining fibroblasts also upregulated transcriptional regulators known to drive ECM target genes (e.g. *Runx1*, *Tgfbr3*, *Egr2*), along with matrix-associated factors and proteases (e.g. *Thbs3*, *Ccn1*, *Mmp2*) related to osteogenic processes **(Fig. 6A)**, suggesting these cells may also play a role in facilitating NGF-induced osteophyte formation. Additionally, Wnt and Smo are also known to regulate endochondral ossification (54–57). To interrogate the transcriptional mediators underlying expression of these NGF-induced genes, we performed a regulon analysis of all lining fibroblast-derived genes significantly upregulated in response to NGF treatment. This analysis predicted one significantly enriched, directly annotated binding motif **(Fig. 6B)**, which is shared among the Sox family of transcription factors. Indeed, in our dataset, lining fibroblasts expressed 4 Sox family genes, *Sox4, Sox5, Sox6*, and *Sox9*, with *Sox5* exhibiting markedly higher expression than all other Sox genes **(Fig. 6C)**. The gene regulatory network for *Sox5* via this binding motif implicates a potential role for Sox5-driven expression of genes related to the fibrosis- and ossification-related pathways activated by NGF **(Fig. 6D)**. Collectively, these findings suggest a key role for lining fibroblasts in driving the pathological effects of NGF in the synovium, with Sox5 identified as the putative transcriptional mediator underpinning the associated transcriptional program.

**Figure 6:**
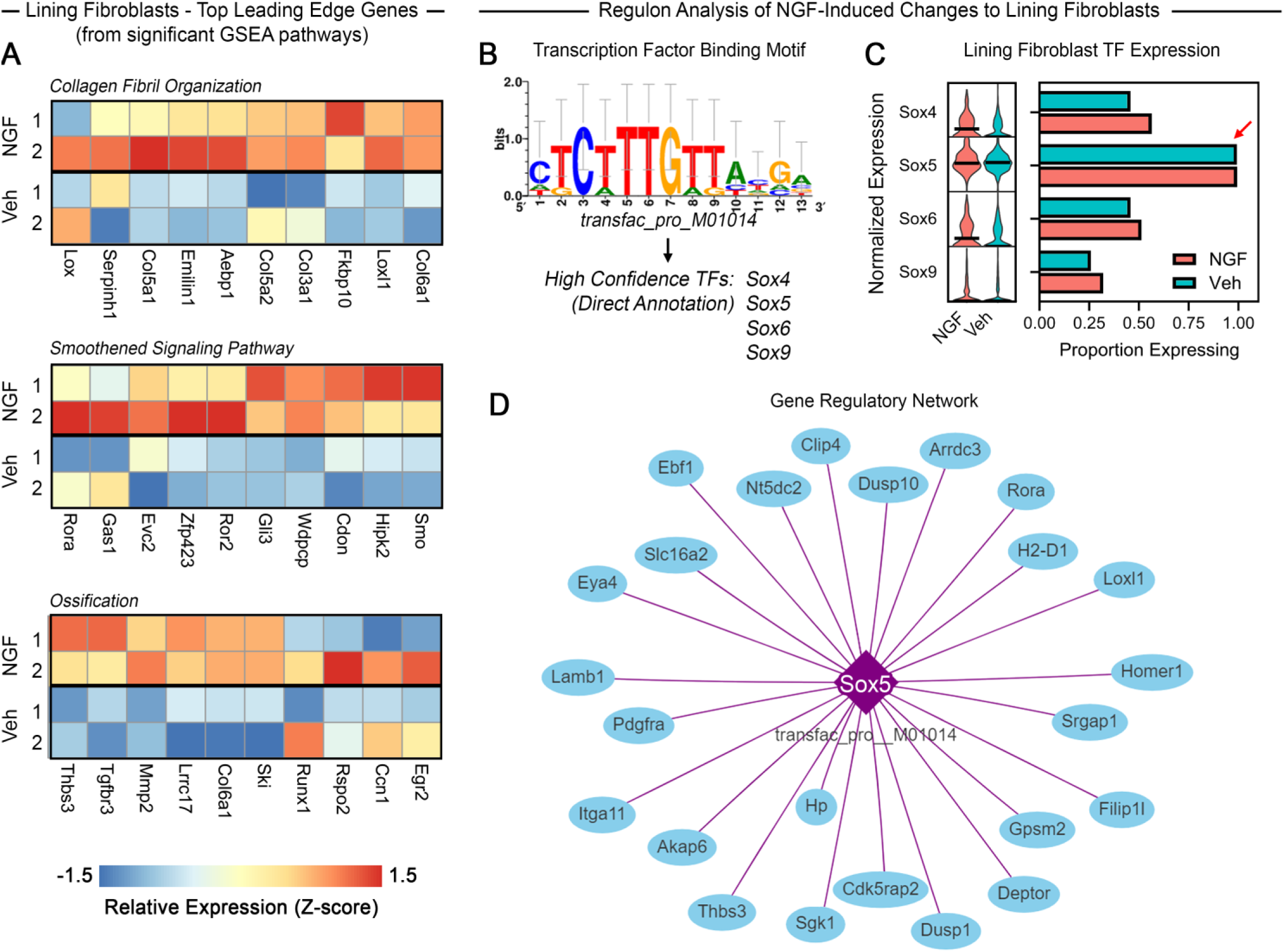
NGF-induced changed in lining fibroblast transcriptome. (A) Heatmap of top gene set enrichment analysis-derived leading-edge genes, demonstrating coordinated upregulation of genes related to collagen fibril organization, smoothened signaling, and ossification; (B) Transcription factor binding motif analysis (RcisTarget) of lining fibroblasts in NGF *vs.* Veh revealed significantly enriched motif transfac_pre_M01014 for the Sox family of transcription factors, predicted by direct annotation; (C) Expression of predicted transcription factors by lining fibroblasts in Veh and NGF groups, demonstrating highest expression of Sox5 (red arrow); (D) Gene regulatory network plot for Sox5 in lining fibroblasts.

### Bulk RNA sequencing of the lumbar dorsal root ganglia

Since NGF is known to be retrogradely transported to the DRG and induce transcriptional changes (5), we sought to describe molecular changes in the DRG that are associated with the phenotypes we observed. We wanted to examine an early time point when mice show knee hyperalgesia but that precedes innervation changes in the joint. Therefore, in this experiment we collected ipsilateral L3-L5 DRGs after 3 IA injections of NGF (500ng) or vehicle for bulk RNAseq. At this time point, we observed 296 genes upregulated and 415 gene downregulated with NGF treatment **(Fig. 7A)**. The top 50 differentially expressed genes are shown in **(Fig. 7B)**, and of these top 50 genes, *Ngfr, Cnr1, F2r, Mmp9, Nptx1, Npy1r* and *Ptgir* have been previously linked to pain (reviewed in (58)), but many had not been previously directly linked to NGF.

**Figure 7:**
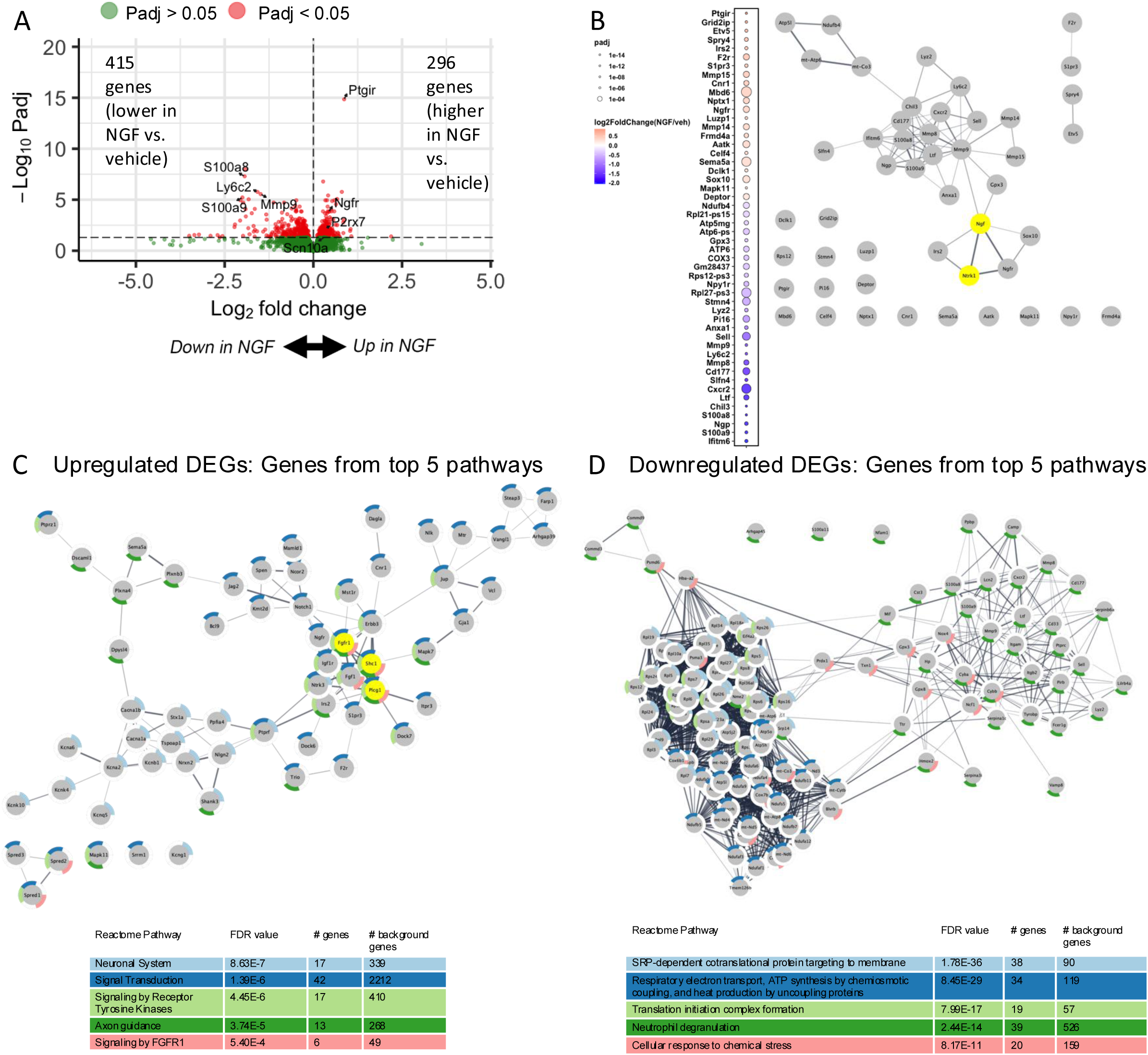
Differential gene expression analysis on bulk RNAseq conducted on DRGs collected after 3 injections of NGF or vehicle. (A) Volcano plot showing all genes that were not filtered out by DESeq2 (15,664). Red indicates genes that met the Padj cut-off < 0.05. No fold change cutoff was applied. Genes of interest are labeled; (B) Top 50 genes by padj, ordered by log2 fold change with the size of the bubble mapped to the adjusted p-value. String network showing known relationships to Ngf and to Ntrk1 (highlighted in yellow); (C) Functional enrichment was performed on the upregulated DEG list using Cytoscape String Enrichment; Interacting genes contributing to the top 5 Reactome pathways are shown using a redundancy cutoff of 0.5. Hub genes are highlighted in yellow; (D) Functional enrichment was performed on the downregulated DEG list using Cytoscape String Enrichment; Interacting genes contributing to the top 5 Reactome pathways shown.

Functional enrichment analysis of the upregulated genes showed the top 5 enriched pathways yielded several pathways related to neuroplasticity: Neuronal system; Signal transduction; Signaling by receptor tyrosine kinases; Axon guidance; and Developmental biology **(Fig. 7C)**. Within the ‘Neuronal system’ pathway, many potassium and calcium ion channel genes were upregulated, as has previously been associated with NGF stimulation of sensory neurons (5). Cytoscape network analysis revealed that *Shc1, Fgfr1, and Plcg1* contribute to all but the ‘Neuronal system’ pathway and represent the genes with the highest degree of connectivity **(Fig. 7C)**. *Shc1* and *Plcg1* have been linked to TrkA signaling (59,60), while NGF has been shown to induce expression of *Fgfr1*(61). All 3 genes are expressed by nociceptors, with *Fgfr1* additionally being expressed by fibroblasts in the DRG, as seen in our scRNAseq database (23). Input of the genes contributing to these top 5 pathways for transcription factor analysis yielded *Prr12*, *Kcnip3*, *Zbtb46*, *Sox10* and *Kmt2a*. Due to the identification of a Sox transcription factor in the synovial analysis as well, it is of interest that *Sox10* was also one of the top 50 genes regulated by NGF in the DRG **(Fig. 7B)**. From our previous scRNAseq data (23), *Sox10* is primarily expressed by Schwann cells and satellite glia in the DRG and has been linked to peripheral nervous system development but not directly to NGF (62).

Functional enrichment analysis of the downregulated genes yielded SRP-dependent cotranslational protein targeting to membrane; Respiratory electron transport, ATP synthesis by chemiosmotic coupling, and heat production by uncoupling proteins; Translation initiation complex formation; Neutrophil degranulation; and Cellular response to chemical stress **(Fig. 7D)**. Genes with the highest degree of connectivity for each of the top 5 downregulated pathways include *Rps3, Atp5h, Rps3, Nme2,* and *Cox6b*, respectively, but none of these genes have previously known links to *Ngf* or *Ntrk1*.

These results provide evidence that intra-articular NGF administration activates a transcriptional program marked by upregulation of genes related to excitability, neuroplasticity and sprouting in the knee-innervating DRG.

## DISCUSSION

In view of the sizeable increase of NGF levels in joints with OA and other arthritides (13,14) and given the limited knowledge of the actions of NGF on joint tissues despite broad expression of its receptors, we investigated the biological effects of intra-articular NGF administration in naïve, adult mouse knee joints. This approach confirmed previous reports of knee swelling and hyperalgesia following IA injection of NGF (22,23). Here, repeated injections of high doses of NGF (500ng) into the knee joint caused knee swelling and hyperalgesia, while knees exposed to a 10-fold lower dose also developed hyperalgesia but not swelling.

While the sensitizing and pain-producing effects of NGF on joints have been extensively reported (22,63–66), to our knowledge no studies have explored the consequences on individual joint tissues of long term exposure to NGF. It has been reported that in all tissues innervated by sensory and sympathetic neurons (such as the skin and internal organs), NGF production is tightly controlled in basal conditions, because alterations of NGF levels can have profound effects on physiology (2). During inflammation, NGF is strongly upregulated, and given the broad expression of its receptors, this can be expected to have profound effects. For example, a significant increase in NGF synthesis in inflamed tissues has been described in patients and animal models of inflammatory diseases (15,19–21,67). Therefore, we opted to repeatedly administer exogenous NGF into naïve murine knee joints and found that this indeed had dramatic effects on the joint. First, we observed that exposure to NGF caused pathological changes in the synovium, with increased subintimal cellularity and fibrosis in addition to lining hyperplasia in the medial and lateral synovium. In experimental OA, an early increase of synovial NGF has been reported to be associated with synovial pathology. First, in the rat monoiodoacetate (MIA) model, NGF protein levels were increased early on (7 days after MIA injection) in proliferating synovium and tissues anterior to the lateral capsule, and NGF levels continued to increase slightly until day 28 (19). Another study reported elevated synovial NGF levels early on in the rat medial meniscectomy model, and this was strongly associated with synovitis in the suprapatellar pouch and medial tibiofemoral compartment (68). Concordant with our findings that NGF-injected mice developed synovial fibrosis, an older *in vitro* study reported that NGF induced proliferation of human fibroblast-like synovial cells in a dose-dependent manner (69).

Synovial pathology induced by NGF was accompanied by robust growth of nociceptors in the medial but not lateral synovium after 4 weeks of repeated injections. Newly sprouted nociceptive fibers were present in deeper synovial layers, away from the lining cells. This is consistent with a recent study by our group, where we used 3D light sheet microscopy to describe innervation changes in response to IA NGF injections into naïve mouse knees, and also showed sprouting in the medial synovium (26). Axonal sprouting is an established biological effect of NGF, and has been reported after endoneurial injection of NGF in rats (70). In the skin, there is also ample evidence to support the notion of NGF-induced nerve growth. Increased NGF content in lesional skin was associated with increased length of epidermal PGP9.5 fibers in human contact eczema (71). In rat skin, exogenous intradermal NGF induced collateral sprouting and treatment with anti-NGF therapy prevented the sprouting (72). In addition, NGF induced sprouting of nerves in aged host rats when ocular transplants were collected from young and old animal donors. The extent of nerve sprouting in the young and old transplants were similar, but sprouting occurred at a slower pace in aged tissue (73). Blockade of NGF using anti-NGF antibodies inhibited sprouting of CGRP+, NF200+ and TH+ fibers and reduced the formation of neuroma like structures in sarcoma injected mice (74).

Two remarkable observations were noted in the neuronal growth response to NGF injections. First, the newly sprouted synovial fibers had a very similar location and distribution to the newly sprouted fibers in the medial synovial sublining we have reported in OA models (30,33), which suggests that nerve sprouting in OA may be mediated by NGF. We are planning to address this in the future by evaluating the effect of NGF blockade on nerve sprouting in OA joints. Secondly, the growth of nerves in response to NGF injections was restricted to the medial synovium, and we saw no sprouting in the lateral synovium in this timeframe. The growth of chondrophytes or pre-osteophytes, was also restricted to the medial compartment even though the whole joint was exposed to NGF. This suggests that the changes in the medial side may be mediated by mechanical loading, since it is known that the medial compartment has higher medial contact load during level walking compared to the lateral side (75). A critical role for mechanical loading in nerve sprouting was also shown in another *in vivo* study, where authors detected elevated NGF mRNA and protein levels in osteoblasts within 1 hour of mechanical loading of the forelimb and inhibition of TrkA signaling reduced load-induced nerve sprouting in the periosteal surface of the ulna (76). These observations support a link between mechanical loading and NGF. Nociceptors express the mechanically gated PIEZO2 channel and the development of mechanical sensitization by NGF has been linked to activity of PIEZO2 expressed by nociceptors (23,77,78). For example, mice where *Piezo2* is knocked out from nociceptors are protected from mechanical sensitization in response to IA injection of NGF (23).

Of note, exogenous NGF did not cause overt cartilage damage in this timespan in spite of the expression of both receptors in chondrocytes (11,15). NGF induced the development of pre-osteophytes, and we observed nerve growth in developing osteophytes. NGF also increased subchondral bone in the medial compartment. An interesting study recently demonstrated the importance of NGF for heterotopic bone formation in a soft tissue injury model, where post-traumatic heterotopic ossification was accompanied by an acute increase in NGF expression. Deletion of *Ngf* or inhibition of TrkA impaired axonal growth into the injured site and reduced the pathologic process of trauma-induced osteochondral differentiation and thus heterotopic bone formation (79). Another study reported robust NGF expression in perichondrial cells close to the ossification center in the developing femur (80). This study showed that inhibition of NGF-TrkA signaling or genetic deletion of NGF from perichondrial osteochondral precursors resulted in disrupted femoral innervation, vascularization, and ossification (79). A further study demonstrated that exogenous NGF increases bone, and that the increased bone is associated with nerve sprouting and this is consistent with our observation of increased BMD and Tb.Th on the medial side (76). Taken together, this suggests that NGF-TrkA signaling plays an important role in both bone formation in normal homeostatic processes and in heterotopic bone formation in response to abnormal conditions. Of note, we did not see increased nerves in subchondral bone channels in response to NGF in this timeframe, which may be due to the fact that NGF could not reach the subchondral bone since the cartilage was intact.

In order to explore molecular mechanisms underpinning the observed biological effects, we performed scRNAseq on synovia collected from injected mice. NGF treatment induced a robust multi-cellular perturbation, particularly within fibroblasts, Schwann cells, and endothelial cells. Synovial lining fibroblasts exhibited the most robust transcriptional response to NGF treatment, with enrichment of pathways (Smo, Wnt) related to fibrosis and matrix deposition, corroborating histological evidence of lining hyperplasia in response to NGF treatment. These findings are consistent with previous reports that lining fibroblasts express NGF receptor p75NTR and expand in response to NGF signaling (81). Lining fibroblasts have also been implicated in osteophyte formation; in response to NGF, these cells were enriched for genes involved in ossification, including upregulation of signaling pathways (e.g. *Smo, Wnt*), transcription factors (e.g. *Runx1*), and matrix-associated factors (e.g. *Thbs3*), which are known to promote endochondral ossification (54–57). The synovial lining is contiguous with the periosteum, and lineage-tracing experiments have previously demonstrated the presence of synovial lining-derived progenitors within developing osteophytes (81), suggesting lining fibroblasts may contribute to osteophyte formation either directly and/or via paracrine signaling with osteochondral lineage cells, and that exogenous NGF injection is sufficient to induce osteophyte formation.

NGF also caused marked alteration of gene expression in the knee innervating DRG. While it has been known that NGF can regulate the gene expression of nociceptor receptors, transmitters, and ion channels (5), this is the first report of transcriptome-wide effects of NGF. Here we focused on persistent NGF effects by collecting DRGs from mice that had been exposed to three IA injections of NGF over the course of 10 days. The bulk RNA sequencing results further supported the neuronal growth promoting effects of NGF. Findings revealed upregulated genes related to axon guidance (*Plxna4, Plxnb3, Sema5a*) and to receptor tyrosine kinase signaling (*Fgfr1, Shc1, Plcg1*), important for axonal growth (82). As we saw in the synovium, genes related to Schwann cell function were also enriched in the DRG (*Sox10*, *Sema5a*, *Erbb3*) (83). NGF injection alters the transcriptional program of Schwann cells, activating multiple pathways related to the support of neuron development including myelination and regulation of neurotransmitter release/synapse function (84–86), suggesting these cells may coordinate with nociceptor afferents to contribute to the neuron sprouting we observed within NGF-treated synovium. Schwann cells play an important role in neuronal development and in facilitating peripheral nerve regeneration following injury, but their role in sprouting in response to non-neuronal tissue damage has been less studied (87). In contrast, we observed downregulation of genes expressed by neutrophils (*S100a8*, *S100a9*, *Mmp9*) and genes related to cell stress in mice injected with NGF compared to vehicle. As neutrophil recruitment to sites of inflammation is a dynamic process, this observation is likely related to the chronic time point chosen for this study.

Several limitations should be discussed. For one, we only evaluated male mice. However, our ongoing studies show NGF causes knee hyperalgesia in female mice as well. We only tested the effect of administering NGF twice a week. The molecular profiling experiments undertaken in this study give us an initial global view on the importance of synovial cells in guiding nerve growth in response to NGF, but future work will be needed to understand the implicated signaling pathways in more detail.

In summary, the introduction of exogenous NGF into healthy young knee joints resulted in profound structural and neuronal remodeling in multiple tissues, mimicking several OA-like changes including synovial fibrosis, increased subintimal cellularity and hyperplasia of the intimal layer of the synovium, development of medial pre-osteophytes, increased thickness of medial subchondral bone and medial synovial nociceptor sprouting. Notably, while NGF is strongly upregulated in OA cartilage (15–18), exogenous IA NGF had no effect on cartilage in this study.

NGF blockade remains an attractive therapeutic strategy for management of chronic OA pain despite the reports of rapidly progressive OA in some patients (9). In view of the profound effects of NGF on all joint tissues, it will be critical to describe the distribution of NGF and its receptors in all joint tissues in great depth, and how they change with age and disease. Such insights will be especially important as they may enable the modulation of NGF signaling to selectively target pain pathways while maintaining its protective functions (88).

## Supporting information

NGF scRNAseq Supplemental Methods

## ABRREVIATIONS

OA: Osteoarthritis
WT: Wildtype
IA: Intra-articular
NGF: Nerve growth factor
ScRNAseq: Single cell RNA sequencing
TrkA: tropomyosin receptor kinase-A
p75NTR: P75 neurotrophin receptor
DRG: Dorsal root ganglia
RPOA: Rapidly progressive osteoarthritis
PAM: Pressure Application Measurement
EDTA: Ethylenediaminetetraacetic acid
H&E: Hematoxylin and eosin
PFA: Paraformaldehyde
ROI: Regions of interest
DMM: Destabilization of medial meniscus
VSMCs: Vascular smooth muscle cells
DE: Differential expression
GSEA: gene-set enrichment analysis
μCT: Micro-computed tomography
BV/TV: Bone volume/total volume
BMD: Bone mineral density
Tb.Th: Mean trabecular thickness
Tb.Sp: Mean trabecular separation

## ACKNOWLEDGEMENTS

Lai Wang.

## FUNDING

This work was supported by National Institutes of Health (NIH)/National Institute of Arthritis and Musculoskeletal and Skin Diseases (NIAMS). AMM was supported by R01AR064251, R01AR060364, and P30AR079206. CRS Grant support VA BLR&D Program (I01BX004912), RR&D Program (I21-RX003854) and NIAMS (R01AR075737). RJM by (R01AR064251), REM by (R01AR077019), and TM by R01AR080035 and R21AR076487. AMO was supported by a T32 Postdoctoral Training in Joint Health (T32AR073157). LL was supported by a Graduate Research Fellowship by the National Science Foundation. The funding sources had no role in the study.

## DECLARATIONS

### Ethics approval and consent to participate

All animal procedures were approved by the Institutional Animal Care and Use Committee at Rush University Medical Center.

### Consent for publication

Not applicable

### Availability of data and materials

Data availability statements should include information on where data supporting the results reported in the article can be found including, where applicable, hyperlinks to publicly archived datasets analysed or generated during the study.

### Competing interests

The following authors declare no conflicts of interest: AMO, MN, BH, SI, JL, LL, LJ, EF, TM, RJM, CRS, REM. AMM received consulting fees from Orion, Roivant, Merck, and Novartis.

### Authors’ contributions

**AO**: Conceptualization, Study design, Data curation, Formal analysis, Investigation, Interpretation, Methodology, Project administration, Writing – original draft, Writing – review & editing. **MN**: Data curation, Formal analysis, Investigation, Interpretation, Methodology, Writing – original draft, Writing – review & editing. **JL, BH, SI, LW, LL, LJ, EF**: Data curation, Methodology, Revised the manuscript. **RJM, CRS, TM**: Conceptualization, Funding acquisition, Resources, Writing – review & editing. **REM**: Conceptualization, Data curation, Formal analysis, Funding acquisition, Investigation, Methodology, Project administration, Resources, Software, Supervision, Validation, Visualization, Writing – original draft, Writing – review & editing. **AMM**: Conceptualization, Formal analysis, Funding acquisition, Investigation, Project administration, Resources, Supervision, Writing – original draft, Writing – review & editing.

## SUPPLEMENTAL FIGURE

**Suppl. Figure 1:**
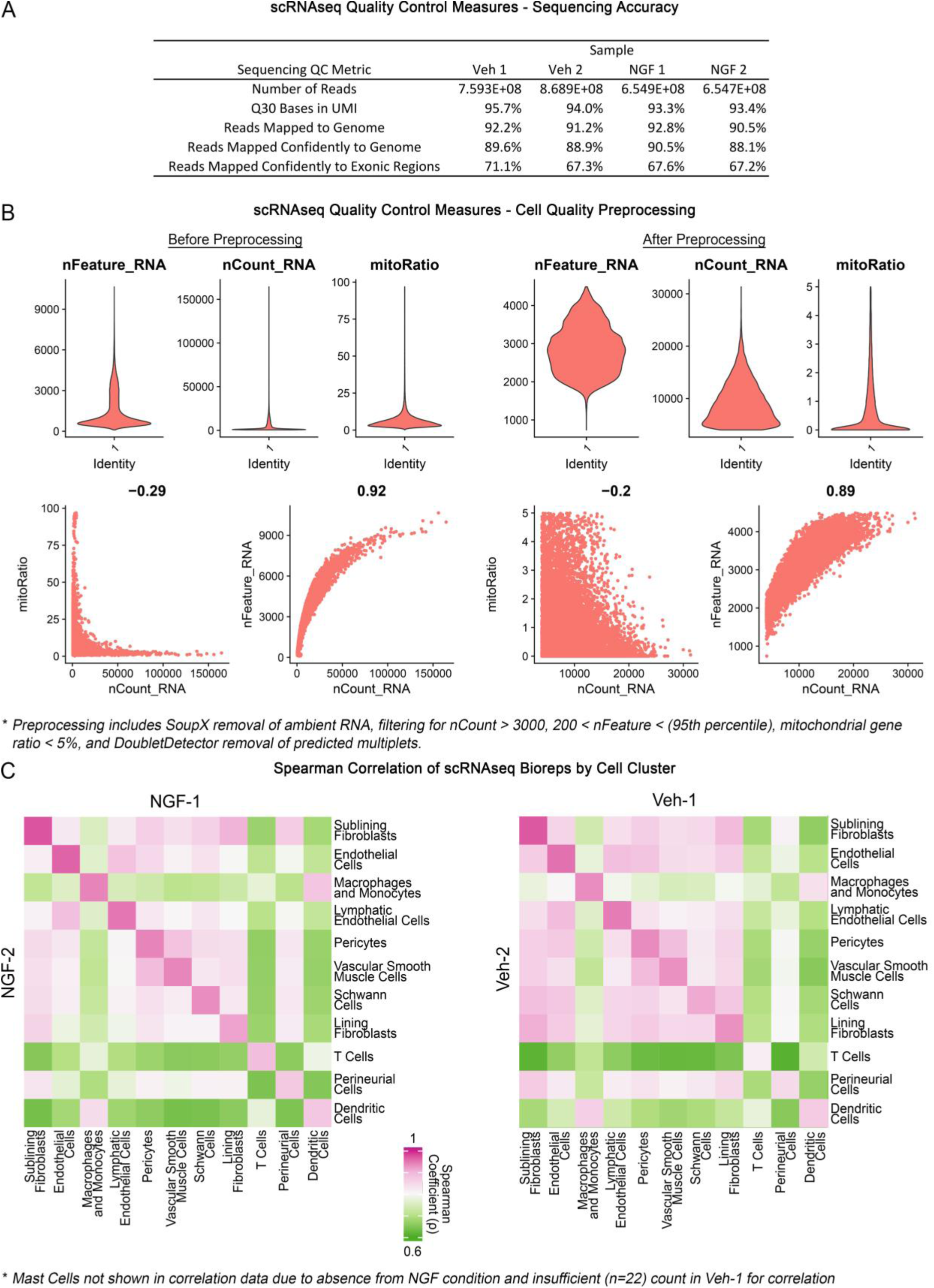
Single cell RNA sequencing quality control. (A) Sequencing accuracy quality control measures; (B) Cell quality metrics before and after preprocessing; (C) Spearman correlation of bio replicates.

**Suppl. Figure 2:**
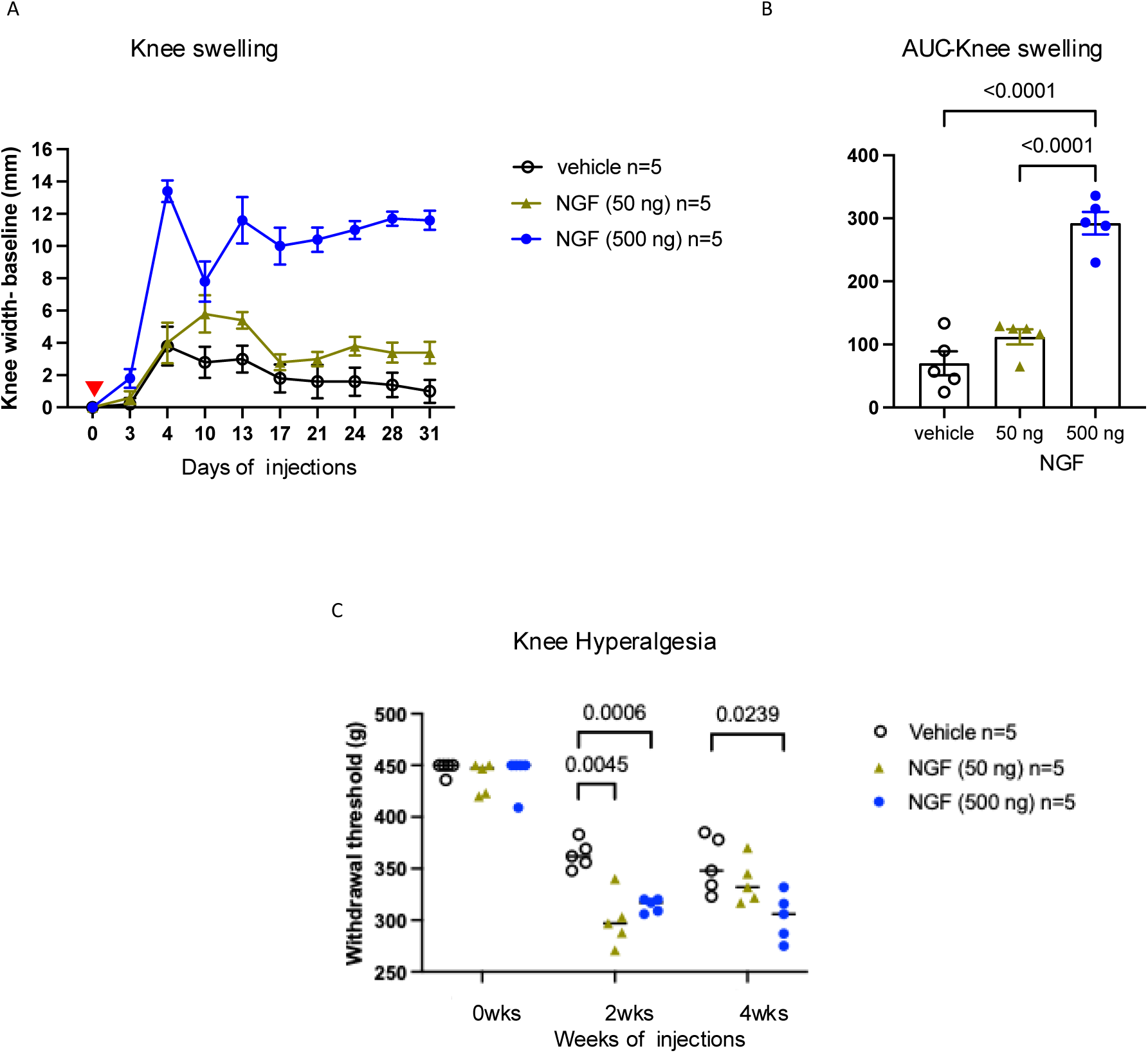
(A) Knee swelling was assessed in wild-type (WT) mice given repeated intra-articular injections of recombinant murine NGF (50 ng or 500 ng, 5 µL, 2x/week) or vehicle (5 µL, 2x/week) for 4 weeks; n=5 mice/group. Red arrowhead indicates the first injection; (B) Knee swelling shown as the area under the curve analysis over the time course. Two-way ANOVA with Sidak’s post-test. One-way ANOVA with Tukey’s post-test; (C) Knee hyperalgesia was assessed for mice given repeated intra-articular injections of recombinant murine NGF (50 ng or 500 ng, 5 µL, 2x/week) or vehicle (5 µL, 2x/week) for 4 weeks at 0-, 2, and 4-week. (A-C) Two-way ANOVA with Dunnett’s post-test.

**Suppl. Figure 3:**
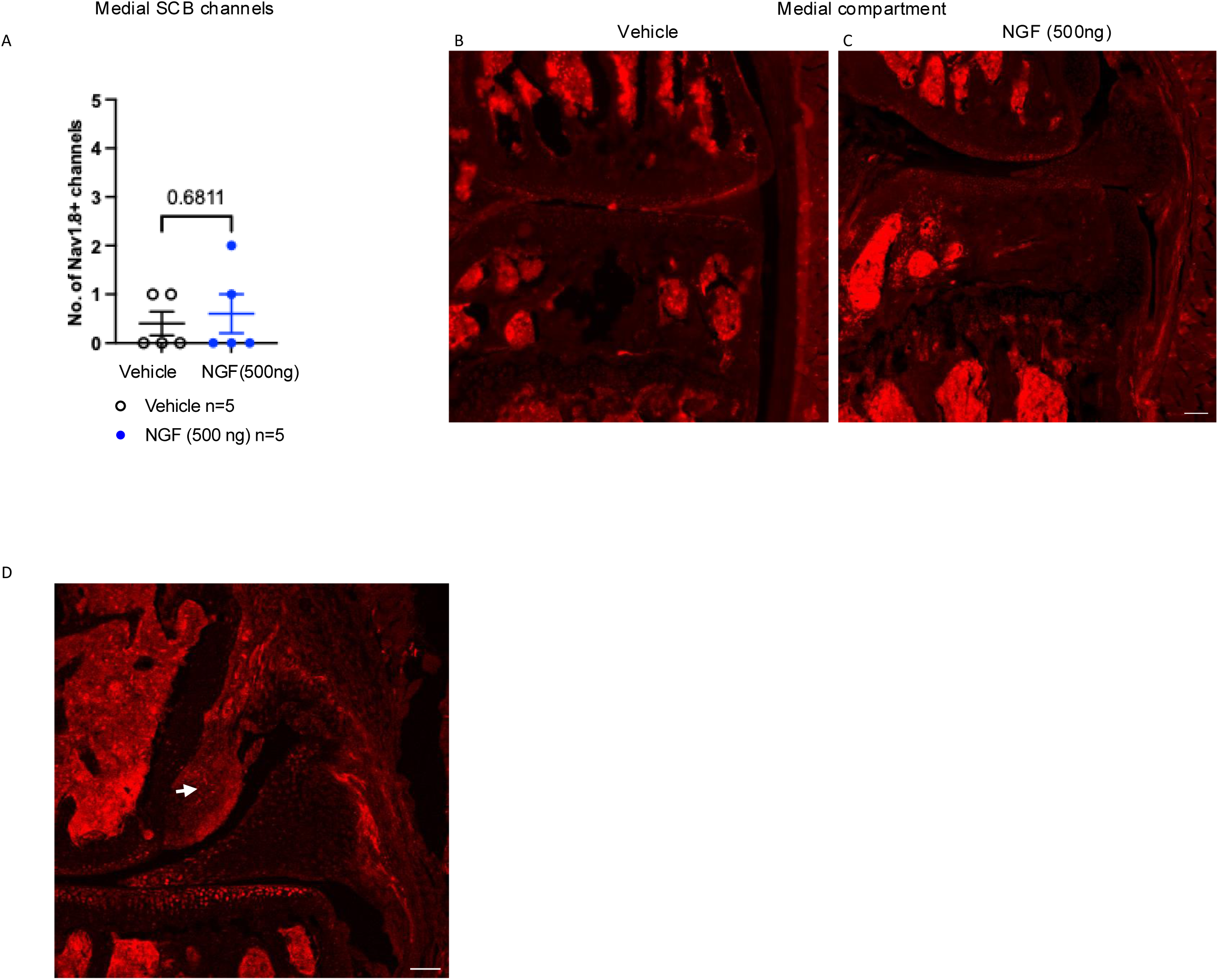
Quantification of Na_V_1.8+ subchondral bone (SCB) channels in the medial compartment of Na_V_1.8-tdTomato mice injected with vehicle (5 µL, 2x/week) or recombinant murine NGF (500 ng, 5 µL, 2x/week) for 4 weeks; n=5 mice/group (A); (B,C) Representative confocal images of the medial compartment of vehicle and NGF treated groups, respectively. Unpaired two-tailed t test; (D) Representative confocal image showing Na_V_1.8+ fibers in pre-osteophytes in white arrow. Scale bar=100µm.

**Suppl. Figure 4:**
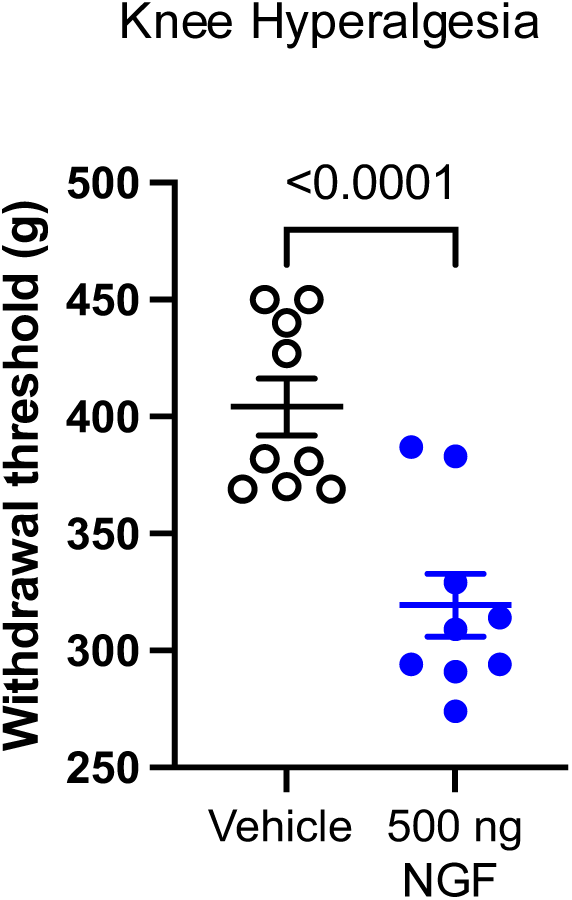
Knee hyperalgesia was assessed at the 4-week time point for mice given repeated intra-articular injections of recombinant murine NGF (500 ng, 5 µL, 2x/week) or vehicle (5 µL, 2x/week) for 4 weeks (Expt 4), and this cohort was used for the synovium scRNAseq experiment (Expt 4); n=9 mice/group were used to measure knee hyperalgesia, however only n=4 randomly selected mice/group were used for scRNA sequencing. Unpaired two-tailed t test.

**Suppl. Figure 5:**
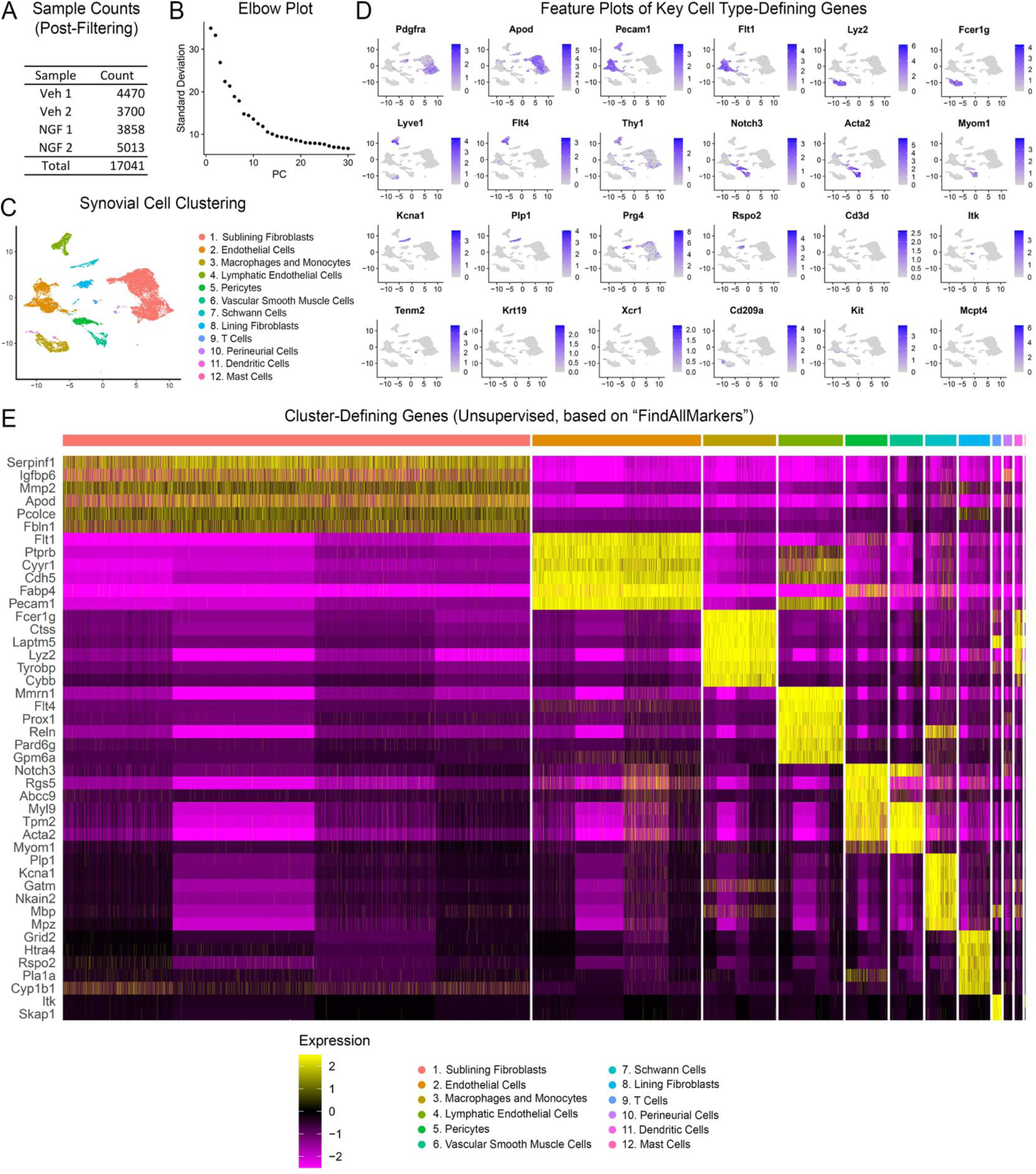
Counts and clustering of synovial single cell RNA sequencing data. (A) Cell counts per sample following pre-processing to exclude low quality cells and debris; (B) Elbow plot from principal component analysis showing contributions of components 1:30 to overall observed variance; (C) UMAP of synovial cell clustering; (D) Feature plots of established cell type gene markers; (E) Heatmap of top cluster-defining genes, based on the “FindAllMarkers” function (pct.1 > 0.7, pct.2 < 0.3, sorted by Padj).

**Suppl. Figure 6:**
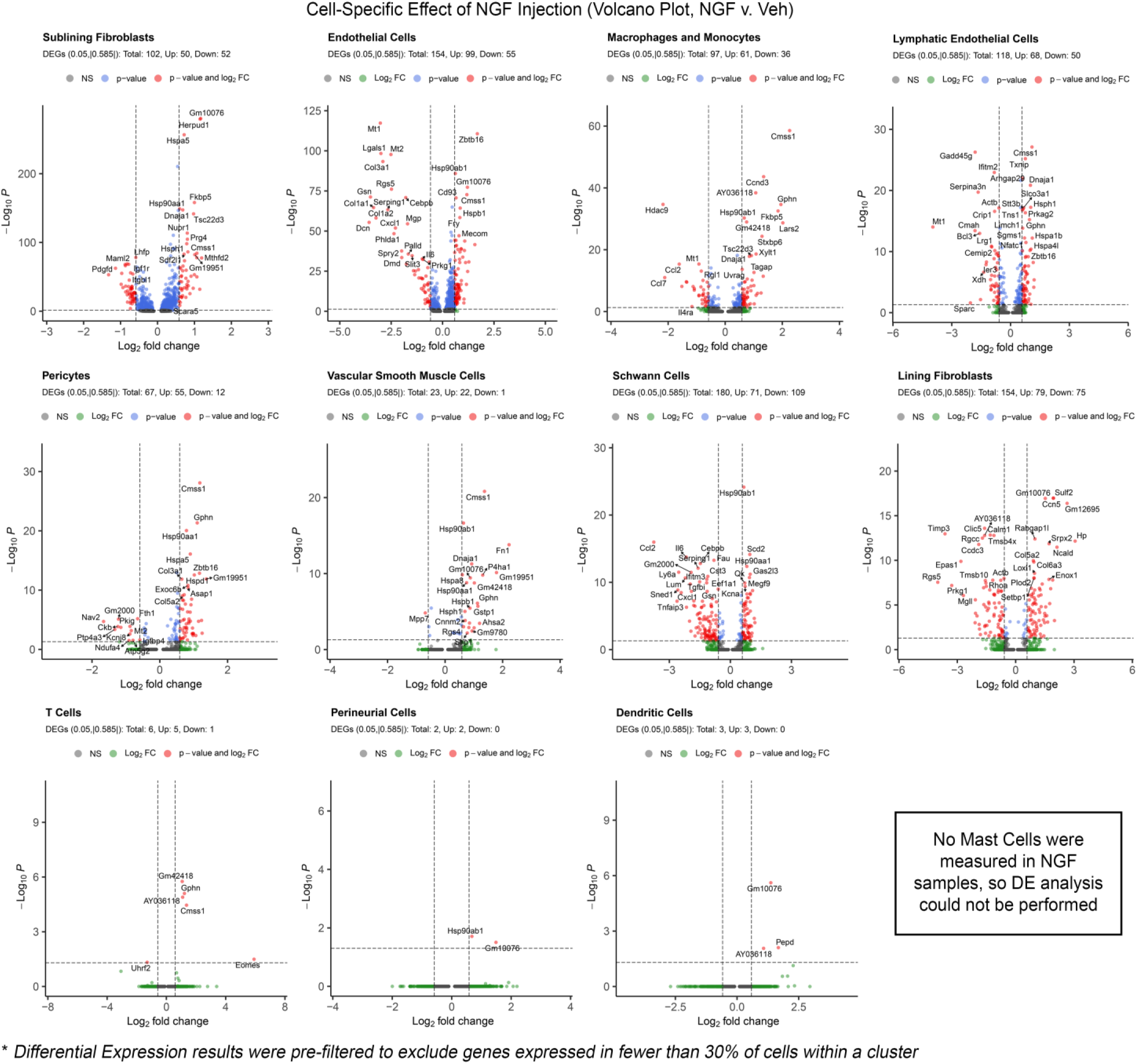
NGF-Induced differential gene expression by cell cluster, shown as volcano plots. Cutoffs at FDR < 0.05 and |log2(fold change)| > 0.585.

**Suppl. Figure 7:**
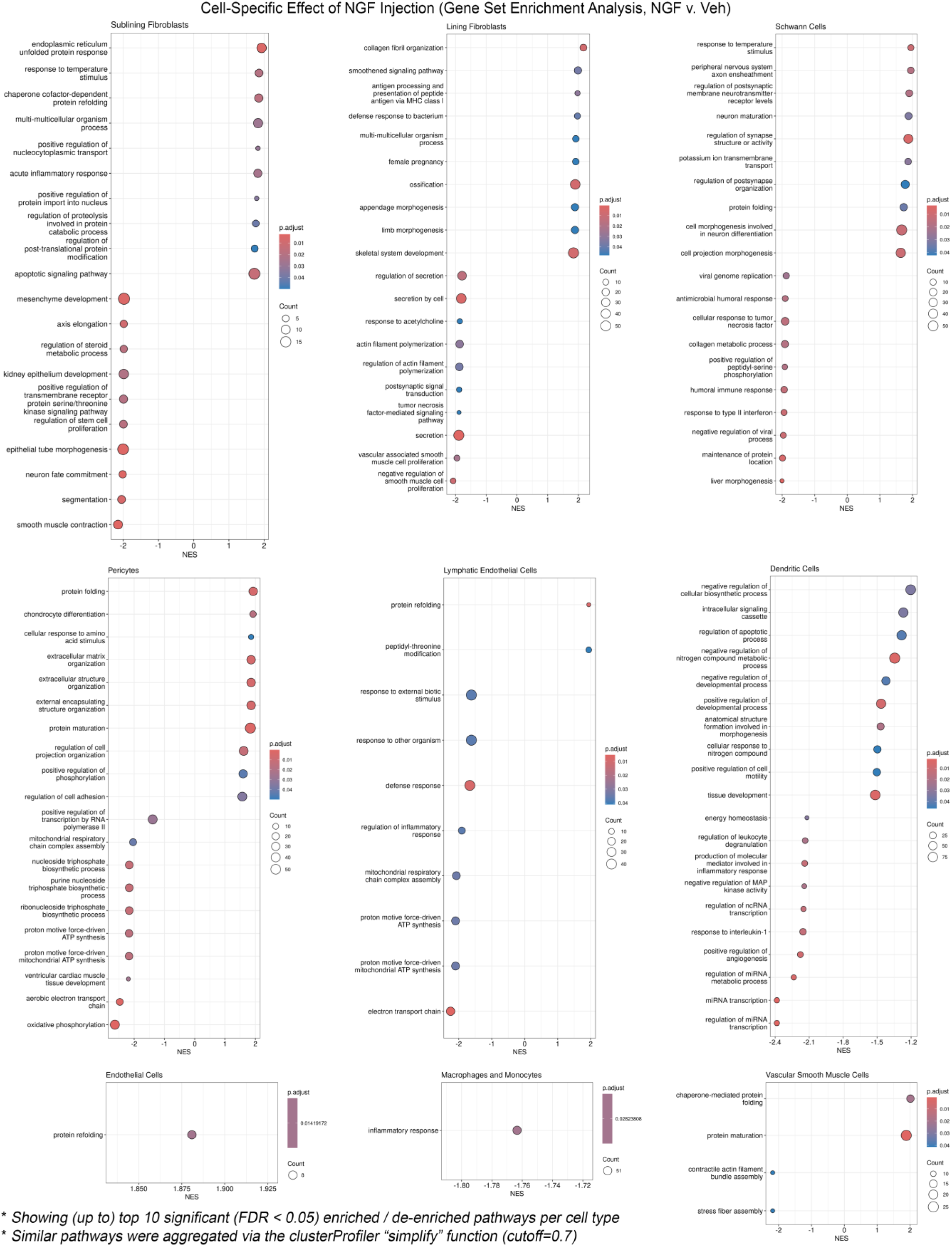
NGF-Induced gene set enrichment by cell cluster, shown as dot plots of up to top 10 significantly enriched and de-enriched terms (FDR < 0.05, sorted by normalized enrichment score (NES). Clusters not shown had no significantly-enriched terms.

