## Supplementary material for "NERVE GROWTH FACTOR IS SUFFICIENT TO CAUSE MULTIPLE OSTEOARTHRITIS-RELEVANT PATHOLOGICAL FEATURES IN NAÏVE MURINE KNEE JOINTS": NGF scRNAseq Supplemental Methods

**Single cell RNA sequencing data analysis methods**

For single cell sequencing, 8 C57Bl/6 mice (*n* =2/sex/condition, age 12 weeks) were randomized to receive 4 weeks of biweekly intra-articular injections of NGF (500 ng) or vehicle control for 4 weeks. At the end of week 4, mice were humanely euthanized via CO2 asphyxia, and ipsilateral synovia were collected and digested to yield a single cell suspension, as previously described [ref Alex ARD]. Cell suspensions were frozen in liquid nitrogen until the time of submission. Single cell suspensions were submitted to the University of Michigan Advanced Genomics Core for barcoding and library prep via the 10X Genomics processing pipeline (Chromium Next GEM Single Cell 3’ Kit v3.1). Two bioreplicates were submitted per condition, with each bioreplicate pooled from one male and one female ipsilateral synovium. Pooled libraries underwent paired-end sequencing (100bp+100bp) on the DNBseq G400 system (MGI Tech). Raw data were aligned to the mm10-2020-A mouse transcriptome using STAR [1] and were preprocessed in Cell Ranger (v7.0.1). Quality control of sequencing indicated high accuracy across all samples (**Suppl. Fig. 1A).**

Analysis of count matrices from Cell Ranger was performed in R (v4.4.1) via the Seurat toolbox [2] (v5.1.0). Counts were preprocessed using SoupX [3] (v1.6.2) to remove ambient RNA contamination. Briefly, raw and filtered feature matrices from cell ranger were imported into Seurat. The filtered matrix was normalized and scaled, and variable features calculated via SCTransform [4] (v0.4.1), dimensional reduction was performed via RunPCA and RunUMAP (dim=1:30), and cells clustered (FindNeighbors, FindClusters) using default settings to provide a rough annotation of distinct cell types, separation of ambient from cell-specific RNA to be tailored to different cell types. A contamination fraction of 10% was specified, ambient RNA was then predicted and corrected for.

SoupX-preprocessed count matrices were then individually loaded into Seurat and pre-filtered. Objects with nCounts < 4000 or nFeature < 200 and were excluded to remove debris, objects with nFeature above the 95th percentile were excluded to remove multiplets, and objects with a mitochondrial gene ratio > 5% were excluded to remove non-viable cells (**Suppl. Fig. 1B)**

To further reduce the presence of multiplets in the dataset, particularly harder to detect homotypic and heterotypic doublets, DoubletFinder [5] (v2.0.4) was utilized. Briefly, the dataset was again normalized and scaled, variable features were computed, and the data underwent dimensionality reduction (dim=1:30) and clustered (resolution=0.3). Parameter pK was set by running paramSweep over PCs 1:30 and selecting the pK which produces the maximum BCmetric. The homotypic doublet proportion estimate was automatically determined via the modelHomotypic, specifying an estimated doublet rate of 5%, and finally doubletFinder was used to predict and exclude multiplets using these settings. This yielded a final population of preprocessed cells for each bioreplicate.

Following filtering and doublet exclusion, datasets were normalized, scaled, and variable features were computed using SCTransform [4] - this scaling step included regression of mitochondrial ratio and cell cycle gene expression to minimize the effects of these confounding variables on the dataset. Dimensional reduction was performed via RunPCA (PCs=30), and individual samples were integrated into a single Seurat object via RPCA integration (IntegrateLayers). Unsupervised cell type clustering was performed via FindNeighbors and FindClusters to determine distinct cell types. Cluster markers were identified using FindAllMarkers, focusing on significantly enriched markers (*P_adj_* < 0.05, using the Benjamini-Hochberg correction for multiple comparisons) which were expressed in at least 70% of the cluster of interest, and fewer than 30% of clusters in other cell types. Spearman correlation confirmed strong correlation between cell types across bioreps (**Suppl. Fig. 1C**).

Differential expression (DE) analysis between NGF and Veh was performed within each cell cluster via FindMarkers, with min.pct > 0.3 to avoid genes minimally-expressed within a cluster. Mitochondrial and ribosomal genes were excluded from downstream analysis. Significantly differentially-expressed genes (DEGs) were defined as those with *P_adj_* < 0.05 and absolute log2 fold change > 0.585 (e.g. |fold change| > 1.5). DE results within each cluster were functionally annotated using gene-set enrichment analysis (GSEA) via the clusterProfiler [6] (v4.12.0) implementation of the fgsea algorithm [7] (v1.30.0). A ranked list of genes was submitted to gseGO using the GO biological processes database, with gene rankings calculated as:

$sign\left( {log}_{2}\left( fold change \right) \right)* -{log}_{10}(Pvalue)$	(1)

Unadjusted *P*-values from DE analysis were used to calculate gene rankings to avoid ties. GSEA results with *P_adj_* < 0.05 were defined as significant. From the preliminary list of significantly enriched pathways, an aggregated list was generated using simplify to aggregate overlapping/similar terms (cutoff=0.7), and these results are reported.

Transcription factor binding motif analysis (i.e. regulon analysis) was performed via RcisTarget [8] (1.24.0). A list of significant DEGs used as input. Motif rankings (mm9, 500 bp upstream) and annotations (motifAnnotations_mgi) were acquired from Bioconductor. Recovery curve AUC was calculated via calcAUC, with aucMaxRank set to the default value of 5% the total number of rankings in the database. Significantly enriched binding motifs were calculated via addMotifAnnotations (nesThreshold=3), and genes associated with each motif were annotated via addSignificantGenes (method=‘iCisTarget’). Significant motifs (NES > 3) with high confidence, directly annotated transcription factors associated with them were focused on. Predicted transcription factors were screened to determine which were measurable within the cluster of interest, with mean expression > 0.05 read counts/cell and max expression > 25 read counts/cell used as filters.
